# A nutrient-responsive hormonal circuit controls energy and water homeostasis in *Drosophila*

**DOI:** 10.1101/2020.07.24.219592

**Authors:** Takashi Koyama, Selim Terhzaz, Muhammad T. Naseem, Stanislav Nagy, Kim Rewitz, Julian A. T. Dow, Shireen A. Davies, Kenneth A. Halberg

## Abstract

The regulation of systemic energy balance involves the coordinated activity of specialized organs, which control nutrient uptake, utilization and storage to promote metabolic homeostasis during environmental challenges. The humoral signals that drive such homeostatic programs are largely unidentified. Here we show that three pairs of central neurons in adult *Drosophila* respond to internal water and nutrient availability by releasing Capa-1 and -2 hormones that signal through the Capa receptor (CapaR) to exert systemic metabolic control. Loss of Capa/CapaR signaling leads to intestinal hypomotility and impaired nutrient absorption, which gradually deplete internal nutrient stores and reduce organismal lifespan. Conversely, hyperactivation of the Capa circuitry stimulates fluid and waste excretion. Furthermore, we demonstrate that Capa/CapaR regulates energy metabolism by modulating the release of the glucagon-like adipokinetic hormone, which governs lipolysis in adipose tissue to stabilize circulating energy levels. Altogether, our results uncover a novel inter-tissue program that plays a central role in coordinating post-prandial responses that are essential to maintain adult viability.

## INTRODUCTION

The ability to maintain osmotic and metabolic homeostasis in response to environmental challenges is a fundamental prerequisite for animal life. Organisms must ensure a balanced equilibrium between nutrient intake, storage and expenditure, which is governed by complex interorgan communication mediated by systemic factors that coordinate the action of specialized organs^1,2^. Remarkably, animals that differ dramatically in life history strategy – like humans and insects – engage similar homeostatic responses, such as promoting nutrient uptake during nutrient-poor states and stimulating waste excretion during nutrient-replete states. While these similarities suggest conserved underlying mechanisms, the physiological adaptations and inter-organ communication networks that drive such changes remain unresolved.

To implement the correct homeostatic program, animals must be able to sense internal nutrient abundances, and to adjust their uptake and utilization of resources accordingly. In mammals, the hypothalamus acts as a central command center for nutrient and water sensing, as it contains neuronal populations that are activated by changes in extracellular sugar concentration or osmolality and release key hormones that initiate compensatory organ activities^3,4^. Similarly, in the fruit fly *Drosophila melanogaster*, discrete populations of neurosecretory cells function as nutrient- or osmosensors, which upon stimulation secrete neurohormones that modulate food intake, energy mobilization, gut peristalsis or renal secretion^5–8^. Both mammals and flies thus regulate organ activities in response to internal state, and in many instances accomplish this regulation by similar mechanisms. However, the hormonal factors involved in coordinating systemic osmoregulation and energy homeostasis, as well as the mechanisms by which the signals are integrated by different organs are incompletely characterized. Understanding these processes is of vital importance as failure in these systems often results in fatal consequences for the organism.

The actions of Capa signaling in insects have so far been restricted to post-embryonic stages, during which the Capa peptides are released into circulation from subsets of neurosecretory cells in the central nervous system (CNS), to activate their receptor – the Capa receptor (CapaR) – on target tissues, including the renal tubules, heart and hyperneural muscles, to control diuretic and myotropic functions^9^. In *Drosophila*, the *Capa* gene encodes a preprohormone that is processed to produce three different peptides, Capa-periviscerokinin-1 and -2 (Capa-PVK-1 and -2) and Capa-pyrokinin-1 (Capa-PK-1)^10^. Strikingly, the Capa precursor is differentially processed within different subsets of neurosecretory cells, with a truncated form of Capa-PK-1 released from the subesophageal ganglion (SEG) neurons, while both Capa-PVKs and -PK are secreted from the ventroabdominal (Va) neurons^11^. The Capa peptides are members of the PRXamide peptide family, which is evolutionarily and functionally related to Neuromedin U (NmU) signaling in vertebrates^12^. In mammals, NmU signaling coordinates key physiological processes including visceral-muscle contractions, gastric acid secretion and insulin release as well as feeding and energy homeostasis, and is therefore an attractive therapeutic target for treating obesity^13^.

Here, we report that Capa-PVK signaling (hereafter just Capa) plays a key role in regulating adult osmotic and metabolic homeostasis in response to environmental conditions. Using several genetic strategies, we provide a comprehensive overview of CapaR expression, which identifies the visceral musculature and neuroendocrine cells of the corpora cardiaca (CC) – producing the glucagon analog adipokinetic hormone (AKH) – as targets of Capa action. Loss of Capa/CapaR signaling reduces intestinal contractility, gut compartmentalization and nutrient absorption, which result in systemic metabolic defects characterized by pronounced hypoglycemia and lipodystrophy. These metabolic effects cause a gradual loss of muscle function due to dysregulated Ca^2+^ homeostasis in skeletal muscles, which impairs feeding behavior and cause premature death. We further show that the Capa Va neurons, but not the SEG neurons, secrete Capa peptides in response to nutrient availability to repress AKH release to restrict energy mobilization from the fat body during nutrient-replete states. We propose that the Capa Va neurons operate as post-ingestive osmo- and nutrient-sensors that coordinate a CNS-gut-renal-fat body signaling module to activate two separate pathways: one to enhance intestinal and renal activities to promote energy and fluid homeostasis, and another to inhibit AKH-mediated energy mobilization to prevent hyperglycemia. Our work thus uncovers an adult-specific inter-tissue program that is essential to maintain osmotic and metabolic homeostasis in *Drosophila* – a program that shows remarkable functional similarity with NmU signaling in mammals.

## RESULTS

### Capa/CapaR signaling is essential for adult fly survival

To gain insight into the physiological actions of Capa/CapaR signaling, we used the binary GAL4/UAS system to silence *CapaR* gene expression using RNAi (Fig. S1a). Remarkably, whereas fly development progressed stereotypically (*i.e.* juvenile growth and developmental time are unaffected; Fig S1b-c), knockdown of *CapaR* expression in cells expressing this receptor using *CapaR^RNAi^* driven by the *CapaR* promoter (*CapaR>*)^12^ resulted in strong mortality shortly after adult emergence. The penetrance of phenotypic effects scaled with temperature due to the thermal flexibility of the expression system^14^ (Fig. 1a; median survival time was 10, 3, and 1 days at 18°C, 25°C, and 30°C, respectively). RNAi specificity was confirmed with two additional independent *CapaR^RNAi^* lines that produced similar phenotypes (Fig. S1d). However, to unambiguously demonstrate a post-developmental role of Capa signaling in sustaining adult survival, we adopted an alternative genetic strategy in which we designed a UAS-inducible tissue-specific CRISPR/Cas9 construct for *CapaR* (Fig. S1a), which we spatio-temporally restricted to the adult stage using the TARGET system^15^. Consistent with the phenotypes observed following *CapaR>CapaR^RNAi^* knockdown, these data showed that adult-specific *CapaR* knockout (*CapaR^ts^>CapaR^KO^*) caused complete fly mortality within 10 days following transfer to the restrictive temperature; crucially the lethality caused by *CapaR* deletion was almost fully rescued in flies additionally carrying a *UAS-CapaR* transgene immune to CRISPR-induced mutation (*CapaR^ts^>CapaR^KO^, CapaR*) (Fig. 1b). These results show that Capa/CapaR signaling is required for adult survival.

**Figure 1.**
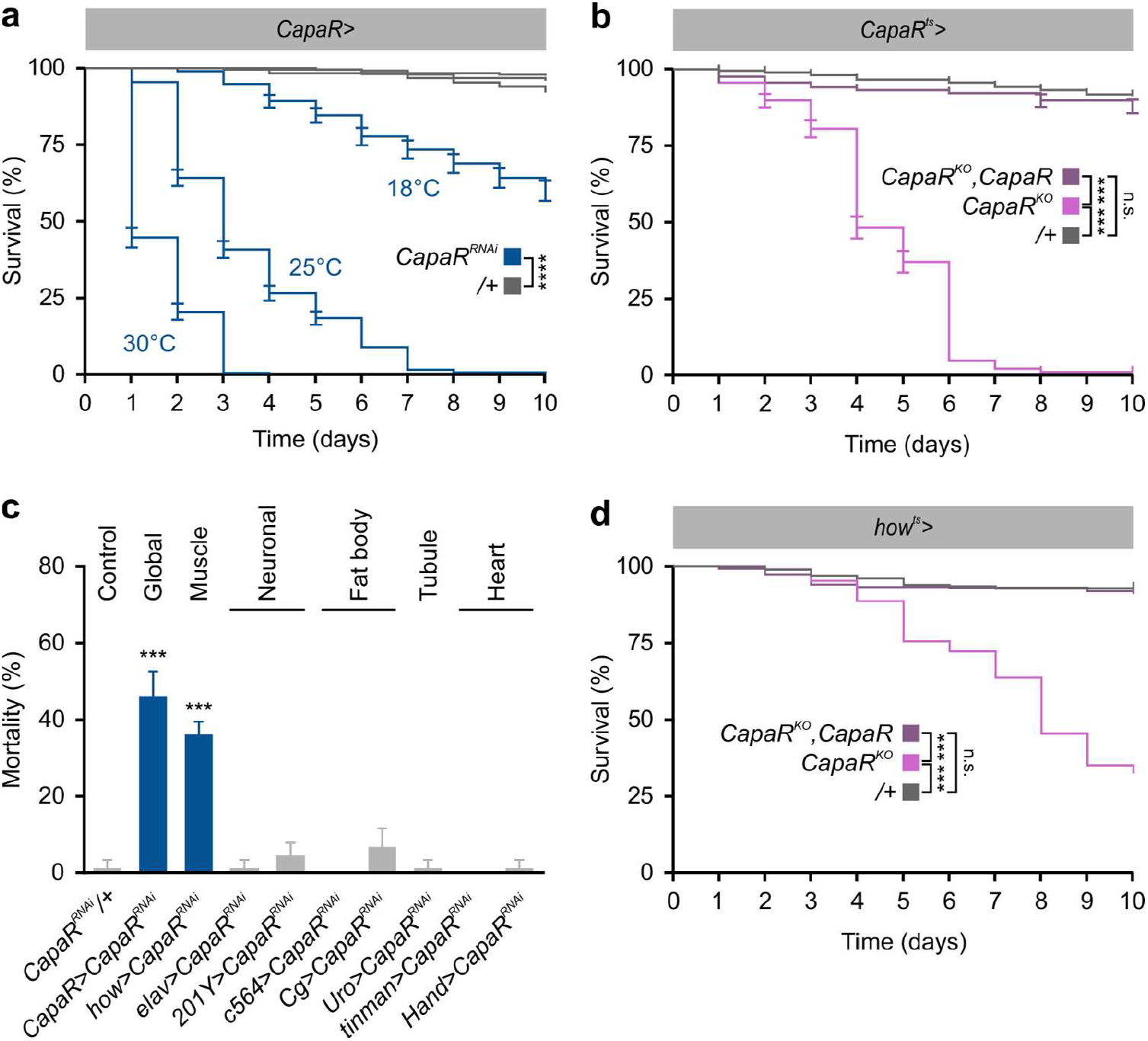
Loss of *CapaR* decreases adult survival. **a.** Global knockdown of *CapaR* (*CapaR> 2xCapaR^RNAi^*) results in strong mortality shortly after adult emergence. Lethality correlates with increased temperature due to the thermal sensitivity of the GAL4/UAS-system. Median survival for male flies was 10, 3, and 1 days at 18°C (n=213), 25°C (n=293) and 30°C (n=232), respectively (log-rank test). **b.** Adult-specific *CapaR* knockout (*CapaR^ts^>CapaR^KO^*) induces significant mortality of male flies (n=498) compared to controls (log-rank test; n=436;). This lethality was fully rescued in flies additionally carrying a *UAS-CapaR* transgene (*CapaR^ts^>CapaR^KO^,CapaR;* n=451). **c.** Knockdown of *CapaR* in whole animals (*CapaR>*) or exclusively in muscles (*how>*), but not in neurons (*elav> or 201Y*), fat body (*Cg*>), Malpighian tubules (*Uro>*) or heart (*tinman> or hand>*), cause significant male fly mortality (one-way-ANOVA; n=60). **d.** Muscle-specific *CapaR* knockout (*how^ts^>CapaR^KO^*) recapitulates the lethality phenotype with knockout flies (n=793) showing significantly higher mortality compared to control (log-rank test; n= 723), and survival is completely rescued in flies co-expressing a receptor transgene (*how^ts^>CapaR^KO^,CapaR*; n=600).

To identify the tissue-specific actions underlying this acute mortality, we selectively downregulated expression of *CapaR* in different tissues. Given our previous work showing that Capa peptides control renal function in *Drosophila* and other insects^16^, we initially explored the possibility that *CapaR* knockdown only in the Malpighian tubules (MTs) – where the receptor is abundantly expressed (Fig. S2a-c) – might impact adult survival. Knocking down *CapaR* expression using the *Uro-GAL4* driver – a GAL4 driver exclusively targeting the principal cells of MTs^17^ – did not cause any significant mortality (Fig. S1e), indicating that the lethality phenotype is uncoupled from Capa-dependent regulation of renal function. Indeed, expression analyses on whole animals showed a much higher reduction of *CapaR* mRNA levels (>90% knockdown) when knocked down in all CapaR-expressing cells (*CapaR>CapaR^RNAi^*) compared to tubule-specific knockdown (*Uro>CapaR^RNAi^;* ◻60% knockdown), suggesting that *CapaR* is expressed in tissues outside the renal tubules (Fig. S1f-g). We therefore assayed the phenotypic effects of *CapaR* knockdown in other major tissues, including the muscles (*how>*), neurons (panneuronal *elav> or* mushroom-body-specific *201Y>*), fat body (*c564> or Cg>*), and heart (*Tinman> or Hand>).* These data revealed that only *how>* – a driver that allows targeted expression in the somatic and visceral musculature (Fig. S1h-i)^18^ – could phenocopy the mortality observed with the *CapaR>* driver (Fig. 1c). Importantly, this mortality phenotype was recapitulated by precise targeting of the somatic CRISPR/Cas9 construct to adult muscles (*how^ts^>CapaR^KO^*), which similar to global knockout (Fig. 1b), caused the majority of flies to die within a ten-day period. Again, this mortality rate was completely rescued to levels approaching that of controls in flies additionally expressing a transgenic *CapaR* receptor in the muscles resistant to the CRISPR effect (*how^ts^>CapaR^KO^,CapaR*) (Fig. 1d). Taken together, these findings link Capa/CapaR signaling to targets outside its well-established role in regulating renal function^12^ and suggests that Capa peptides coordinate additional processes in the musculature that are essential to sustain adult survival.

### Capa signaling targets several tissues

To identify the specific cellular targets of Capa signaling, we analyzed CapaR expression using a combination of genetic and molecular reporters. As expected, *CapaR-GAL4* was found to induce strong reporter activity (*UAS-mCD8::GFP*) in both the anterior and posterior MTs (Fig. S2a-c); however, we detected additional prominent and consistent GFP fluorescence in the thoracic skeletal muscles, brain as well as circular visceral muscles confined to the proventriculus, midgut and rectal regions of the intestine (Fig. 2a; Fig. S2a). The accurate reporting of endogenous *CapaR* localization was verified using a custom-synthesized CapaR antibody^12^, which showed substantial overlap between GFP fluorescence and anti-CapaR immunoreactivity (Fig. 2a). These findings were further corroborated using fluorescently tagged Capa-1 peptide (Capa-1-F) in combination with a recently developed ligand-receptor binding assay^16^; fluorophore labelling did not affect the biological efficacy of the peptide (Fig. S2d-g). Using this approach, we observed specific and displaceable Capa-1-F binding to *CapaR-GAL4-driven* GFP-positive visceral muscles and enteroendocrine cells (Fig. 2b-c) of the adult digestive tract, which is consistent with single-cell transcriptomic data reported by Flygut-seq^19^ showing *CapaR* expression almost exclusive in these two cell types of the gut (Fig. S2h).

**Figure 2.**
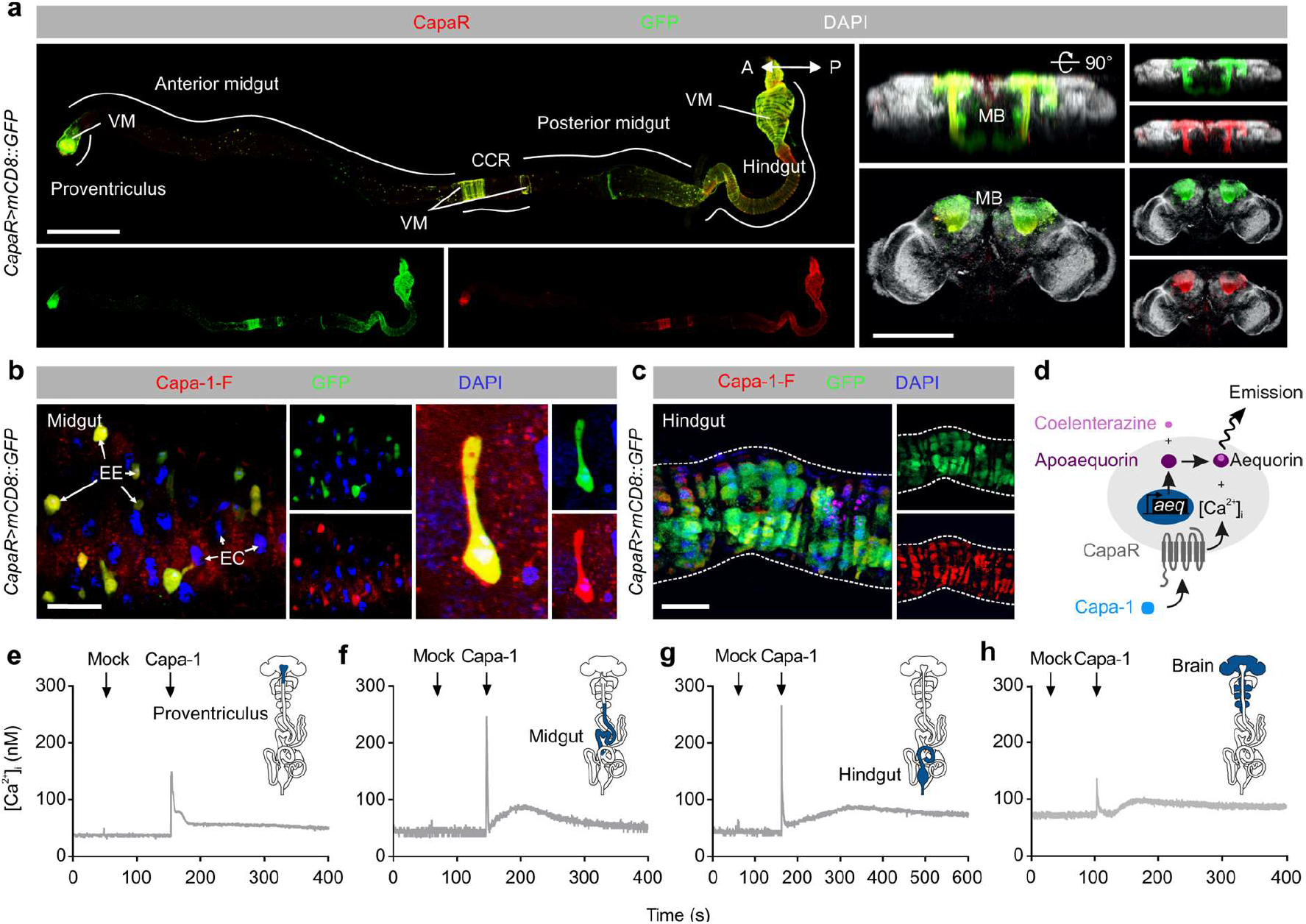
Global CapaR receptor mapping in adult *Drosophila*. **a.** Co-immunolabeling of the CNS and the intestine from *CapaR>GFP* flies with a custom generated CapaR antibody. Consistently, we observed an overlap between GFP (green) and CapaR (red) immunoreactivity in the mushroom body and neuronal somata of the CNS, as well as in the visceral muscles and enteroendocrine cells restricted to distinct portions of the gut. **b-c.** Application of fluorophore-labelled Capa-1 (Capa-1-F, red) on the intestine shows specific and displaceable binding to *CapaR>GFP* (green) positive enteroendocrine cells and visceral muscles. **d.** Principle of *in vivo* aequorin luminescence-based functional calcium assay in *Drosophila* tissues. **e-h.** Real-time changes in intracellular Ca^2+^ concentration of tissues natively dissected from flies expressing a *UAS-apoaequorin* transgene under the control of *CapaR>.* Cytosolic [Ca^2+^]^i^ levels (nM) in **e.** proventriculus, **f.** midgut, **g.** hindgut and **h.** brain upon Capa-1 stimulation (10^−7^ M). MB, mushroom body; EE, enteroendocrine cells; EC, enterocytes; CCR, copper-cell region.

Finally, to functionally validate our receptor mapping data, we expressed an aequorin-based bioluminescent Ca^2+^-sensor (*CapaR>aeq::GFP*) to detect robust changes in cytosolic [Ca^2+^]_i_ following Capa stimulation of acutely dissected tissues (Fig. 2d). Consistent with previous findings, Capa-1 induced a stereotypic, biphasic rise in [Ca^2+^]_i_, comprising a rapid primary peak followed by a slower secondary rise in the renal tubules (Fig. S2i)^12^ Yet more strikingly, we also observed a prominent cytosolic calcium response in immuno-positive gut regions such as the proventriculus, midgut and hindgut/rectum, as well as in the brain (Fig. 2e-h), and to a lesser extent in the legs (Fig S2j); by contrast, we were unable to detect any Ca^2+^ response in the salivary glands or gonads (Fig. S2k-l). Taken together, our combined approaches provide a comprehensive overview of CapaR expression in adult *Drosophila* unmasking several novel target tissues of Capa neuropeptides including the skeletal and visceral muscles.

### Four pairs of Capa neurons mediate brain to peripheral organ communication

To determine the neurons responsible for the Capa-mediated effects on adult viability, and to enable their subsequent genetic manipulation, we characterized a new GAL4 driver line, *Trp-GAL4 (Trp>*), that when crossed to *mCD8::GFP* produced almost complete overlap in immunoreactivity with an anti-Capa precursor antibody^10^ in the adult CNS, hereby demonstrating unparalleled specificity and potency to the *Capa^+^* neurons (Fig. 3). Indeed, knocking down *Capa* expression (*Trp>Capa^RNAi^*), or completely ablating the Capa-producing neurons via targeted expression of the proapoptotic gene *rpr (Trp>UAS-rpr*) recapitulated the observed mortality phenotype of *CapaR* manipulations, with flies dying shortly after eclosion (Fig. S1j). Performing a detailed neuroanatomical analysis of *Trp>UAS-mCD8::GFP* flies, we found consistent GFP expression in one pair of neuroendocrine cells in the SEG, whose dendritic projections innervate the AKH-producing CC cells (APCs) and extend along the esophagus to innervate the proventriculus (Fig. 3a-c; Fig. S3a,c). In addition, we observed reporter activity in the three pairs of Va neurons; the two anterior pairs send axons to a neurohemal plexus in the dorsal neural sheath, while the posterior-most pair join the median abdominal nerve (MAN; Fig. 3d; Fig. S3b), from which their axons project to form neurohemal release sites. Although direct innervation of the intestine by neurons exiting the CNS through the MAN or the presence of Capa^+^ enteric neurons would offer an attractive model for Capa-mediated changes in intestinal physiology, we were unable to observe such connections. In sum, our data reveals neuronal innervation of the CC and proventriculus by the SEG neurons, and the systemic release of Capa peptides into circulation by the Va neurons to activate peripheral target tissues.

**Figure 3.**
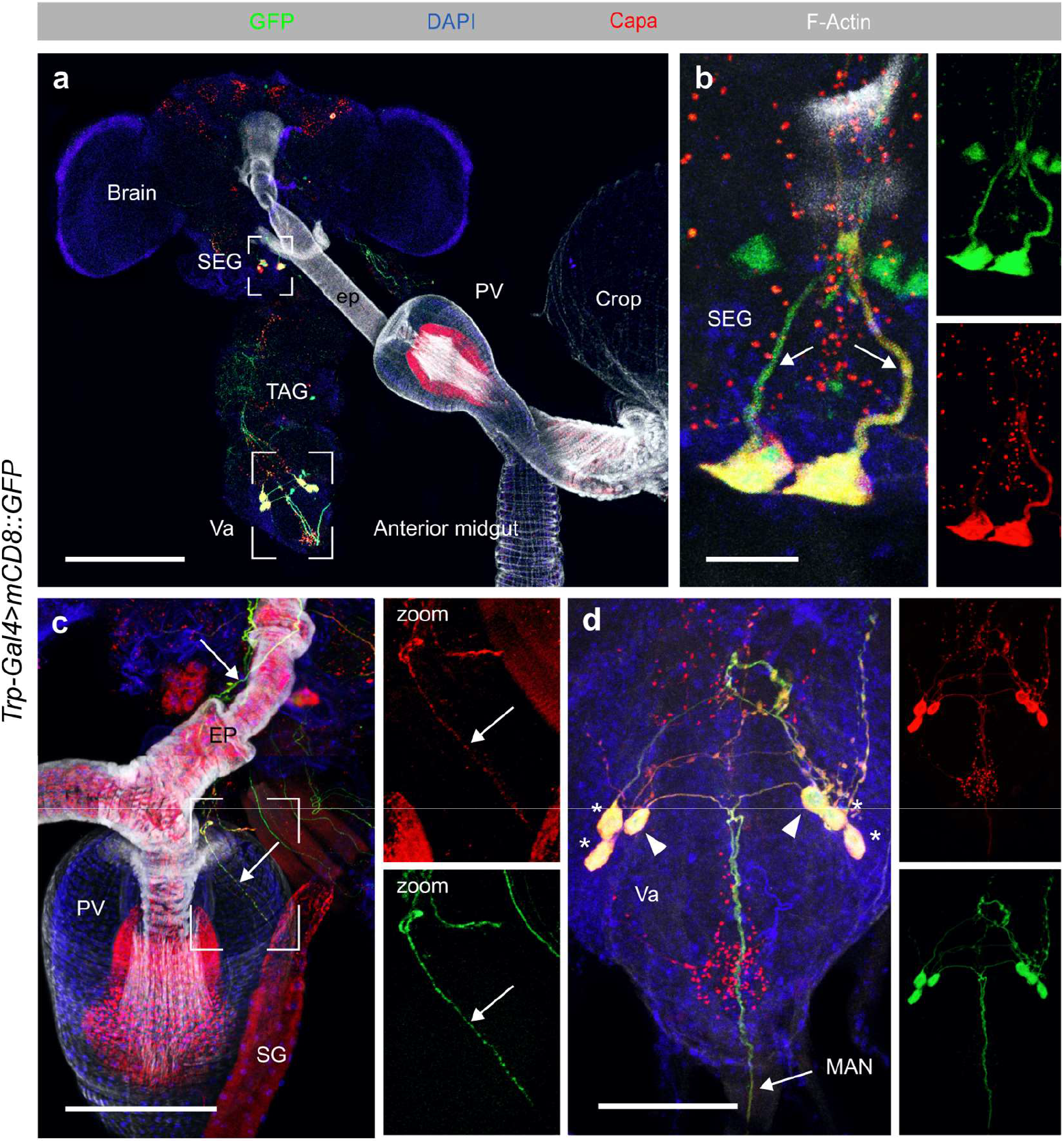
Anatomy of Capa-producing neurons in adult *Drosophila*. **a-d.** The CNS and associated tissues showing *Trp>UAS-mCD8::GFP* (green) counterstained with an anti-Capa antibody (red), F-Actin (white) and DAPI (blue) in maximum projected z-stack confocal images **a.** GFP expression and anti-Capa immunoreactivity co-localize in four pairs of Capa^+^ neurons; one pair in the SEG and three pairs in the TAG. Scale bar, 200 μm. **b.** The SEG neurons extend dorsally. Scale bar, 20 μm. **c.** The SEG neurons also extend along the esophagus to innervate the proventriculus. Scale bar, 50 μm. **d.** The two anterior pairs (*) of Va neurons send axons to the dorsal neural sheath, while the posterior pair (triangles) projects into the median abdominal nerve and form neurohemal release sites. Scale bar, 50 μm. EP, esophagus; MAN, median abdominal nerve; PV, proventriculus; SEG, subesophageal ganglion; SG, salivary gland; TAG, thoracicoabdominal ganglion; Va, ventroabdominal.

### Capa/CapaR signaling regulates gut motility as well as fluid and waste excretion

The expression of CapaR in visceral muscles confined to portions of the adult intestine containing valves and sphincters (proventriculus, posterior midgut and rectal pad), suggests modulation of intestinal transit by Capa-dependent control of smooth muscle cell activity. Such a role is consistent with the known functions of mammalian Neuromedin U receptor 2 (NmU-R2, a functional homologue of CapaR^12^), which has been shown to promote gastrointestinal motility in mammals^20^. Using an established *ex vivo* gut-contraction assay^16^, we quantified the spontaneous contraction frequency of the intestine before and after Capa-1 or Capa-2 application to gut preparations. We found that both Capa peptides – at a concentration previously determined to cause maximum receptor occupation^12^ – stimulate gut contraction frequency approximately 2-3-fold compared to the artificial hemolymph control saline (Fig. 4a-b). Conversely, the Capa-induced activation of gut motility was abolished after *CapaR* knockdown (*how^ts^>CapaR^RNAi^*), indicating that the increase in gut-contraction frequency following Capa-1/-2 application depends on CapaR function in intestinal muscle cells (Fig 4a-b). CapaR is a Gq-protein coupled receptor in which receptor occupancy results in elevation of intracellular Ca^2+^-levels. We therefore used the genetically encoded Ca^2+^-indicator *UAS-GCaMP6s* to directly visualize Ca^2+^-dynamics in CapaR-expressing circular muscle cells during Capa-1 (10^−7^M) stimulation (Fig. 4c). These data show that Capa-1 application induces prominent intracellular Ca^2+^-transients in muscle cells supporting the notion that Capa/CapaR controls visceral muscle activity.

**Figure 4.**
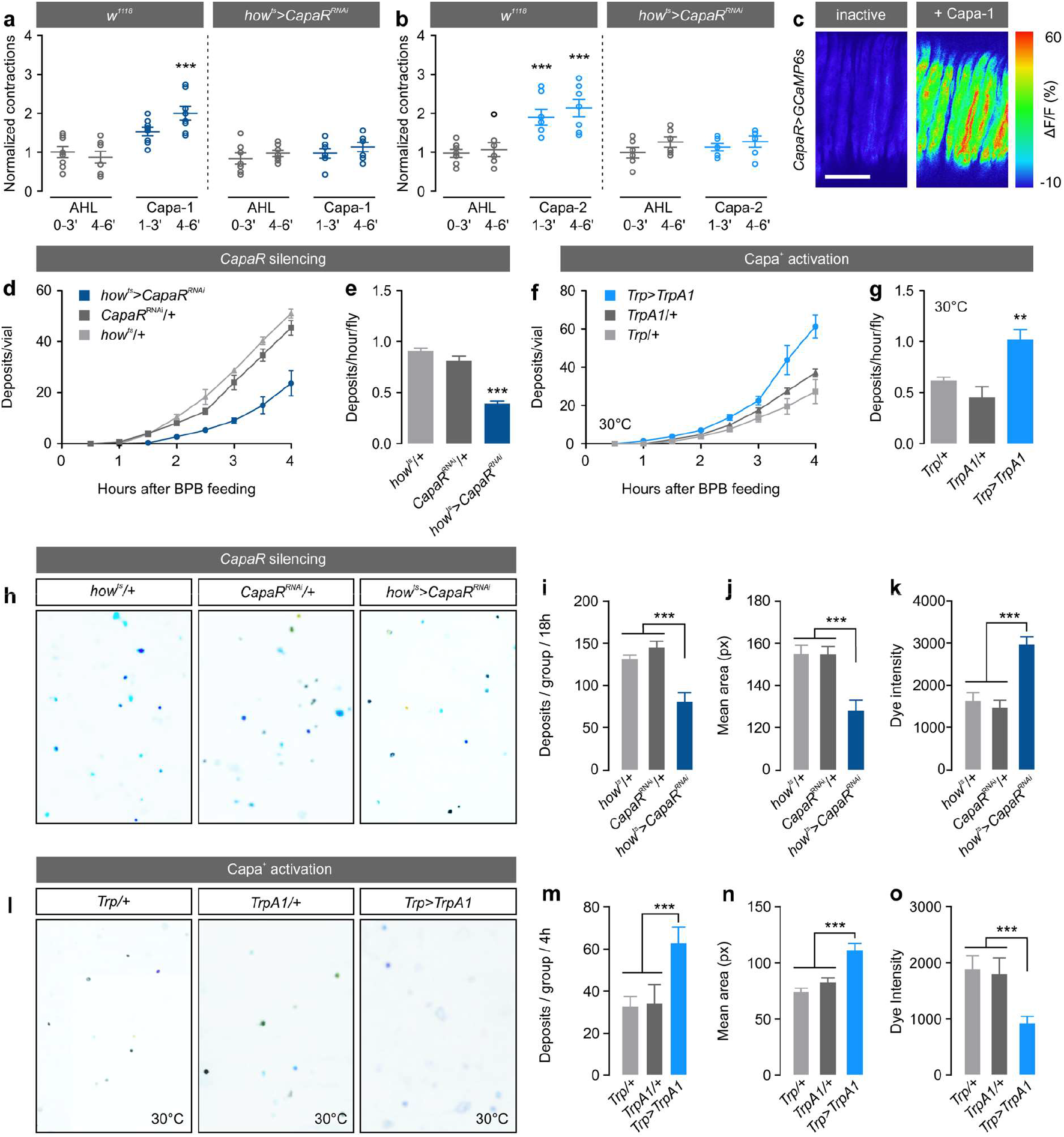
Capa modulates intestinal contractions and excretory physiology. **a-b.** Spontaneous gut contractions from control and muscle-specific *CapaR* knockdown flies (*how^ts^> CapaR^RNAi^*) in response to AHL and AHL containing **a.** Capa-1 or **b.** Capa-2 at a concentration of 10^−7^ M. Grey values indicate exposures to AHL alone, whereas colored values refer to exposures to AHL + Capa-1 or Capa-2, respectively. ΔContractions were calculated by normalizing the number of contractions during Capa-1/-2 application over those in the control (one-way-ANOVA; n=6-8). **c.** Inactive and activated circular visceral muscles in guts from flies carrying *UAS-GCaMP6s* driven by *CapaR>* following Capa-1 stimulation. **d.** Accumulated number of deposits, and e. the number of deposits/hour/fly in muscle-specific *CapaR* silenced flies (*how^ts^> CapaR^RNAi^*) compared to parental controls (one-way-ANOVA; n=3 vials with 20 flies per vial). **f.** Accumulated number of deposits, **g.** and the number of deposits/hour/fly in flies with artificial activation of the Capa^+^ neurons by ectopic expression of the heat-sensitive TrpA1 channel (*Trp>TrpA1*) compared to control flies at 30°C (one-way-ANOVA; n=3 vials with 20 flies per vial). **h, l.** Representative fecal output profiles from the different genetic backgrounds fed on BPB-labelled food. Muscle-specific silencing of Capa signaling (*how^ts^> CapaR^RNAi^*) results in **i.** fewer, **j.** smaller, and **k.** more concentrated BPB-labelled deposits compared to the parental controls (one-way-ANOVA; n=17-19 Petri dishes with 10 flies per dish). **m.** Conversely, artificial overactivation of Capa-producing neurons using a *UAS-TrpA1* transgene induced excretion of more, **n.** larger, and **o.** lighter deposits compared to flies carrying each transgene alone (one-way-ANOVA; n=5-7 Petri dishes with 10 flies per dish).

To explore the physiological significance of this effect, we next asked if the observed impairment of gut motility caused by reduced Capa/CapaR signaling affects intestinal transit time *in vivo*. In flies exposed to food supplemented with the pH-sensitive dye Bromophenol blue (BPB), we observed a significant delay in gut transit in knockdown animals (*how^ts^> CapaR^RNAt^*) compared to parental controls, as evidenced by the delayed appearance and significant decrease in BPB-labeled waste deposits produced over time (Fig. 4d-e). In contrast, genetic overactivation of the Capa-producing neurons using the heat-activated transient receptor potential A1 cation channel TRPA1 (*Trp> TrpA1*) led to a significantly shorter intestinal transit time when incubated at 30°C (Fig. 4f-g). Thus, Capa-1/-2 neurohormones communicate with the gut to modulate intestinal motility and are necessary and sufficient for the control of gut transit in adult *Drosophila.*

To further explore the functional significance of *CapaR* silencing on gut physiology *in vivo*, we adopted an approach that provides quantitative readouts of intestinal emptying rate and fluid content based on the semi-automated analysis of fly excreta^21,22^. These experiments showed that visceral muscle-specific *CapaR* knockdown significantly affects *Drosophila* defecation behavior compared to the parental controls, as these flies produced fewer, smaller, and more concentrated excreta (Fig. 4h-k). These changes in fecal output are likely caused by prolonged contact with intestinal contents associated with gut hypomotility. Consistent with this model, artificial activation of the Capa neurons (*Trp>TrpA1*) led to the production of lighter (less concentrated), larger and more abundant deposits compared to flies carrying either transgene alone, presumably due to the combined effects of intestinal hypermotility and increased diuresis (Fig. 4l-o). Interestingly, limiting the time available for fluid reabsorption in the gut has a larger impact on systemic fluid balance than stimulating renal secretion, as adult-specific expression of a membrane-tethered version of the diuretic hormone DH44 (*Uro>tDh44*) – which allows cell-autonomous activation of its cognate receptor and thus constitutively activate tubule secretion^23^ – failed to fully recapitulate this diarrhea-like phenotype. Instead, the deposits produced by these flies were more numerous, but not larger or more dilute than those of controls (Fig. S4b), which is consistent with the role of the rectum in regulating the final fluid content of excreta^24^. Together, these results show that Capa peptides direct the actions of the CNS-gut axis by modulating intestinal contractility, which is important not only for appropriate gut transit, but also directly impacts the homeostatic regulation of water balance by affecting fluid content of excreta.

### Capa/CapaR regulates gut functions essential to maintain metabolic homeostasis

Next, we asked if these changes in gut physiology affect the metabolic status of the fly. Indeed, in mammals gut hypomotility has been shown to compromise both intestinal acid-base balance, nutrient absorption and organismal energy status^25^. Consistent with these observations as well as our current findings, we observed frequent gut distension as well as loss of acidity in the copper-cell region (CCR) of midguts from Capa/CapaR deficient flies (Fig. 5a); a region functionally analogous to the mammalian stomach^26^. We therefore tested if these gut defects affect nutrient breakdown and absorption by the gut, by analyzing the elemental composition of deposits from flies lacking *CapaR* function in the visceral muscles using scanning electron microscopy coupled with wave-dispersive X-ray analysis. Interestingly, we detected significantly higher levels of carbon and nitrogen (the main elemental components of sugars and amino acids) in the excreta of *CapaR* knockdown flies compared to the control (Fig. 5b-c), suggesting a potential disruption of digestive and/or absorptive functions of the gut. To test this directly, we quantified triacidglyceride (TAG) and glucose levels in the excreta from *how^ts^>CapaR^KO^* flies, which revealed that adult-specific *CapaR* knockout in visceral muscles leads to significantly higher levels of residual undigested nutrients compared to controls (Fig. 5d). Importantly, these gut defects – including the loss of CCR acidity – were fully rescued by co-expressing a UAS-CapaR construct (*how^ts^>CapaR^KO^,CapaR*) (Fig. 5a,d). To further test this model, we exposed *how^ts^>CapaR^KO^* flies to more nutrient-dense foods in order to promote energy-uptake and assessed the impact of different nutrient concentrations on animal viability. These data showed that higher nutrient concentrations significantly prolonged fly survival in a dose-dependent manner, whereas both control and rescue animals were unaffected (Fig. 5e). Together, our results suggest that *CapaR* elimination in intestinal myocytes affects the digestive and absorptive functions of the gut, which has acute effects on organismal lifespan

Reduced nutrient absorption, however, could also be a compensatory reaction to excess food intake. This idea is consistent with the role of NmU signaling in regulating food consumption in rodent models^27,28^. To test this hypothesis, we quantified the duration of food contact as an indirect measure of food intake, in addition to directly measuring food consumption. These data revealed that *CapaR* knockdown flies spent significantly less time in contact with food (Fig. 5f) and that *CapaR* knockout animals reduced their overall dietary intake; an effect that could be reversed by overexpression of CapaR in muscles (Fig. 5g). We also observed similar effects upon targeting the four pairs of Capa^+^ neurons by Capa knockdown (*Trp>Capa*^*RNAi*^; Fig. 5h), indicating that Capa/CapaR signaling play a role in regulating feeding behavior in adult flies. The combined effects of reduced energy intake and impaired intestinal absorption strongly suggest that these flies show severe metabolic defects. We therefore assessed the internal nutritional status of these flies, by quantifying internal carbohydrate levels (glucose and trehalose) as well as whole-body energy stores (glycogen and TAG) at distinct points in time following visceral muscle-specific *CapaR* elimination (*how^ts^>CapaR^KO^*). As expected, these data revealed a gradual decrease in internal energy stores in *CapaR* knockout animals, perhaps due to increasingly poor feeding and nutrient absorption, as revealed by their progressive hypoglycemic and lipodystrophic phenotypes compared to both control and rescue flies (Fig. 5i-l). Altogether, our data suggest that impairment of Capa/CapaR signaling in muscles impairs food intake and cause defects in the digestive and absorptive functions of the gut, which critically impact the metabolism and lifespan of adult flies.

**Figure 5.**
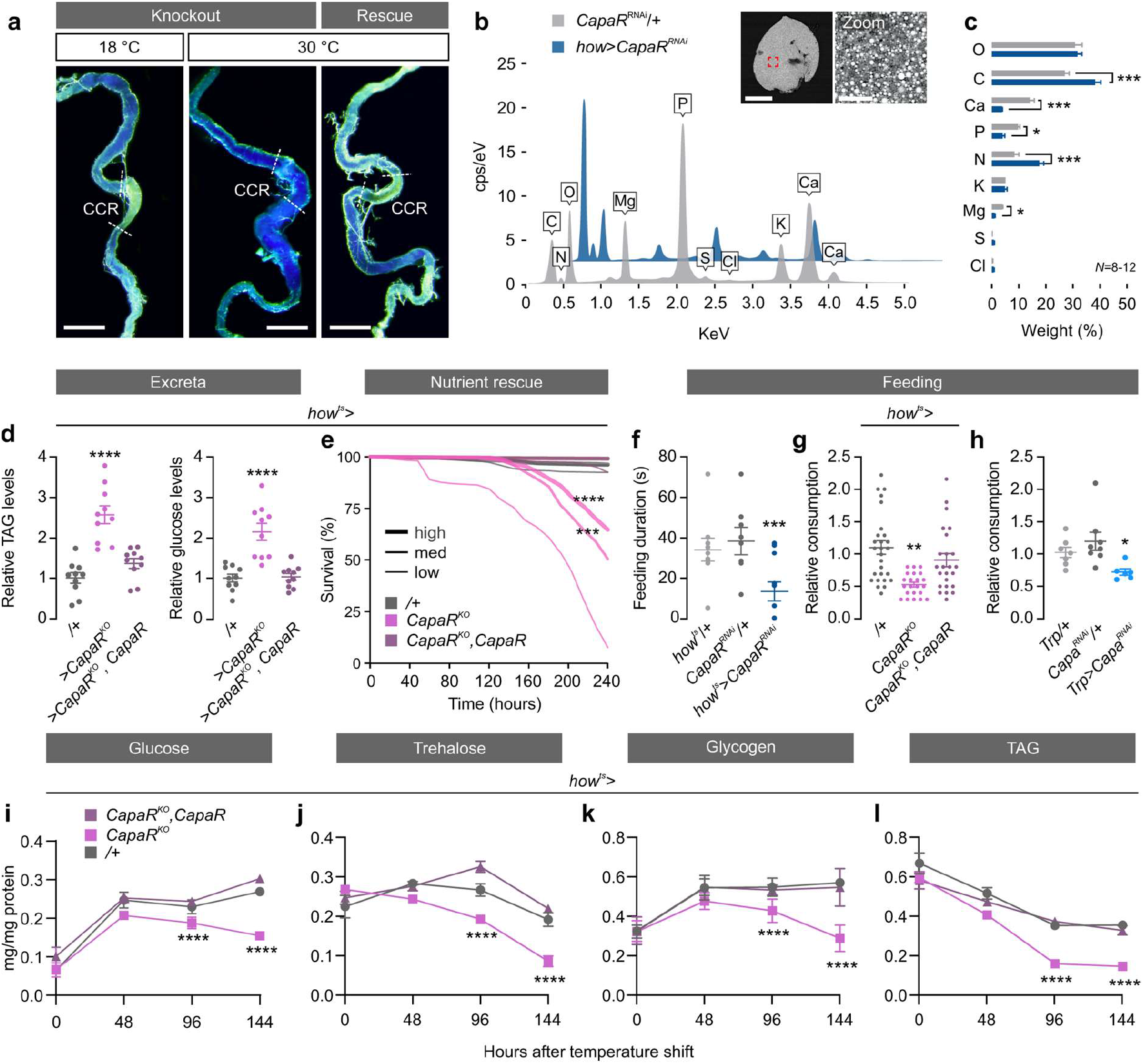
*CapaR* deletion in visceral muscles causes systemic metabolic imbalance. **a.** Dissected intestinal tracts from flies fed with food supplemented with the pH indicator bromophenol blue. A complete loss of acidity in the copper-cell region (CCR) of the midgut was observed, as evident from the absence of yellowish color, exclusively in *how^ts^>CapaR^KO^* flies kept at the restrictive temperature (30°C). This was not detected in animals kept at the permissive temperature (18°C) or in rescue flies kept at 30°C additionally carrying a *UAS-CapaR* transgene (*how^ts^>CapaR^KO^,CapaR*). **b.** Elemental composition and **c.** percentage of weight element in the excreta from visceral muscle-specific *CapaR*-silenced flies (*how>CapaR^RNAi^*) using scanning electron microscopy coupled with wave-dispersive X-ray analyses (unpaired *t*-test; n=8-12). **d.** Relative amounts of undigested triacylglyceride (TAG) and glucose levels in excreta from control (/+), knockout (*how^ts^>CapaR^KO^*) and rescue (*how^ts^>CapaR^KO,^CapaR*) flies raised on standard medium (one-way-ANOVA; n=10). **e.** Flies kept on media with low, medium and high nutrient concentrations show that higher nutrient concentrations reduce mortality in *how^ts^>CapaR^KO^* (low, n=325; med, n=323; high, n=325) flies, with both control (low, n=272; med, n=271; high, n=273) and rescue (low, n=251; med, n=250; high, n=253) flies unaffected by the different nutrient dense diets (log-rank test). **f.** Feeding duration was significantly decreased in *CapaR* knockdown animal (*how^ts^>CapaR^RNAi^*) compared to parental controls. **g.** Relative food intake in *CapaR* deficient (*how^ts^>CapaR^KO^*) flies (n=8) and **h.** *Capa* silenced (*Trp>Capa ^RNAi^*) flies (one-way ANOVA; n=6-8). **i-l.** Quantification of **i.** glucose **j.** trehalose, **k.** glycogen and **l.** TAG levels following 48, 96 and 144 hours of adult-specific transgene activation (one-way ANOVA; n=10).

### Impaired Capa/CapaR signaling reduces muscle performance and adult locomotion

In flies, as well as mammals, changes in systemic energy balance induces the release of hormonal factors that modulate behavioral programs and locomotor activity^29–31^. For example, central administration of NmU peptides increases gross-locomotor activity in mammals^28^. We therefore asked whether Capa/CapaR impairment induces similar responses in locomotor activity, by examining baseline locomotion of adult flies using the *Drosophila* Activity Monitor System. These data revealed that muscle-specific elimination of *CapaR* (*how^ts^>CapaR^KO^*) induced a gradual decrease in adult locomotion over the course of the 4-day experimental period, as evidenced by the significant reductions in both morning and evening activity peaks relative to parental controls (Fig. 6a-b). This observation contrasts with the stereotypic increase in locomotion exhibited by energy-deprived wild-type animals, a behavior linked to food foraging^30^. Conversely, flies with rescued Capa signaling in muscles (*how^ts^>CapaR^KO^, CapaR*) exhibited a pronounced hyperactivity compared to control, perhaps owing to increased Capa/CapaR signaling flux in the musculature of these animals (Fig. 6a-b). To further evaluate the functions of Capa/CapaR signaling in muscles, we videotracked individual flies with either knockdown of *CapaR* in muscles (*how^ts^>CapaR^RNAi^*) or in neurons (*Trp>Capa^RNAi^*) and quantified the total distance traveled as well as maximal response velocity of these animals. Our data confirmed that both muscle-specific elimination of *CapaR* as well as *Capa* depletion in Capa^+^ neurons cause significant reductions in total walking distance and maximal response velocity compared to controls (Fig. 6c-d). These data suggests that CapaR function in muscles is necessary for general motor activity in flies, and that silencing Capa signaling in muscles compromises both short-term and long-term locomotor activity.

**Figure 6.**
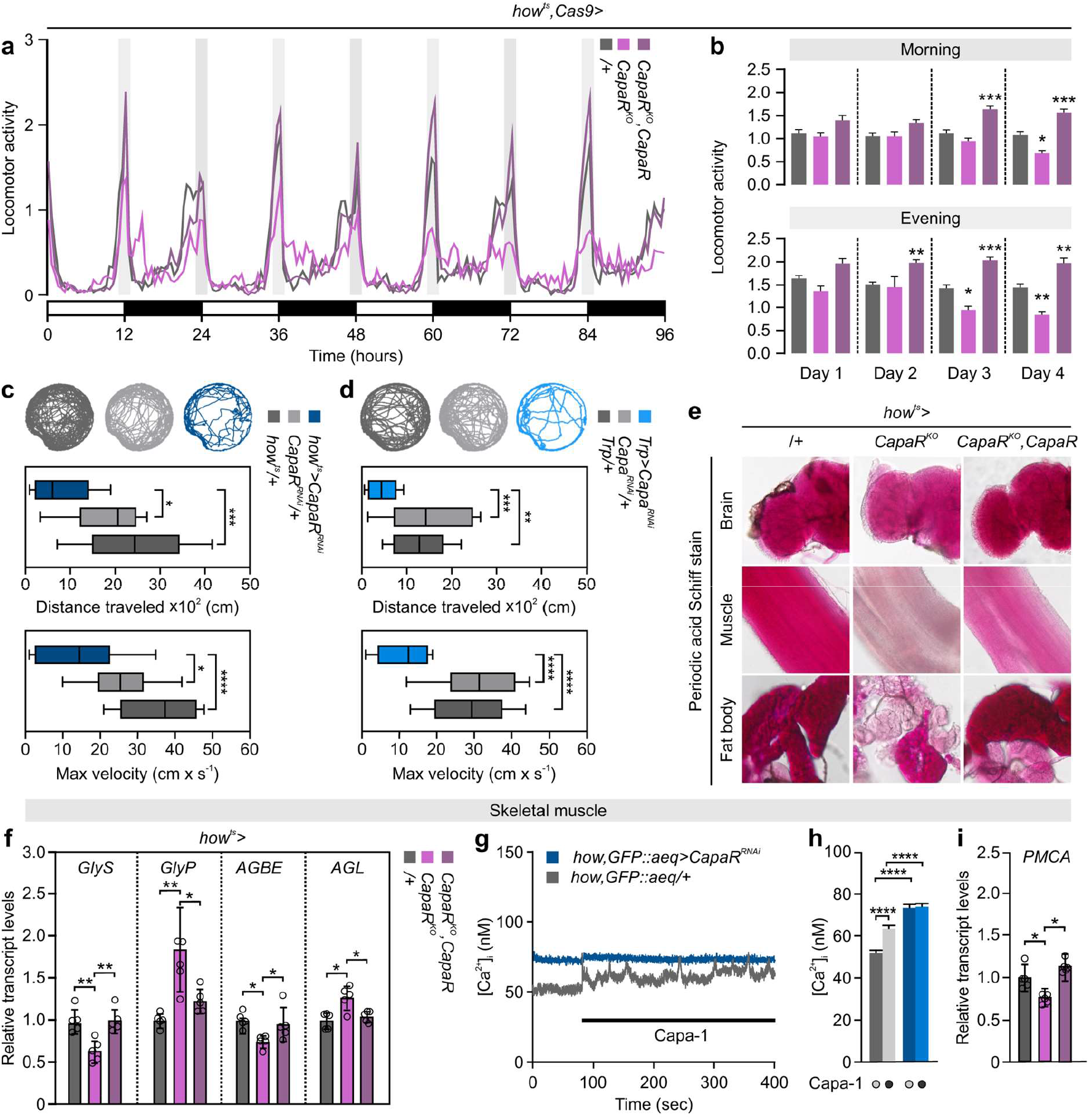
Muscle-specific *CapaR* elimination induces locomotor defects. **a.** Locomotor activity of individual control (/+; n=32), muscle-specific *CapaR* knockout (*how^ts^>CapaR*^*KO*^; n=20) and rescue (*how^ts^>CapaR^KO^, CapaR*; n=32) flies exposed to 12-hour:12-hour, light-dark (LD) cycles for 96 h. The experiment was repeated twice with the same results. **b.** Stereotypic morning and evening activity peaks measured over the experimental period (oneway ANOVA). **c.-d.** Representative activity traces of video-tracked individual flies with targeted *CapaR* silencing in muscles (n=10-16; *how^ts^>CapaR^RNAi^*) or *Capa* knockdown in Capa^+^ neurons (n=10-16; *Trp>Capa^RNAi^*) relative to control flies, including quantifications of distance traveled and max velocity (one-way ANOVA). **e.** Periodic Acid-Schiff (PAS) staining in the brain, muscle, and fat body showed depleted carbohydrate stores in these tissues of *how^ts^>CapaR^KO^* flies relative to control, with partial rescue in flies additionally carrying a *CapaR* transgene (*how^ts^>CapaR^KO^,CapaR*). **f.** Transcript levels of *GlyS, GlyP, AGBE* and *AGL* relative to *RpL32* in skeletal muscles (thorax samples) from muscle-specific *CapaR* knockout (*how^ts^>CapaR^KO^*) and rescue (*how^ts^>CapaR^KO^, CapaR*) flies. Data are expressed as mean fold change compared to control ± s.e.m. (one-way ANOVA; n=5). **g.** Stereotypic cytosolic calcium traces in muscles expressing the aequorin-based bioluminescent Ca^2+^-indicator (*how,GFP::aeq*, grey line), and the effect of attenuating *CapaR* expression (*how,GFP::aeq>CapaR^RNAi^*, blue line) in muscles stimulated with 10^−7^ M of Capa-1 peptide. **h.** Bar graphs of basal and stimulated cytosolic calcium levels in *how;GFP::aequorin>CapaR^RNAi^* and *how;GFP::aequorin/+* control samples Data are expressed as mean [Ca^2+^] (nM) ± s.e.m. (one-way ANOVA; n=3). **i.** Transcript abundance of *PMCA* compared to *RpL32* expression in muscles from control (+/), knockout (*how^ts^>CapaR^KO^*) and rescue (*how^ts^>CapaR^KO^,CapaR*) flies (one-way ANOVA; n=5).

We next explored the underlying cause of this reduced activity. Capa peptides have been shown to possess myomodulatory effects in a range of insects^32–34^, raising the possibility that Capa signaling may directly regulate skeletal muscle contractility in *Drosophila.* We therefore eliminated Capa signaling exclusively in the flight muscles by using the *Act88F-GAL4* driver, hereby isolating the physiological effects produced by skeletal muscles. These data showed that eliminating CapaR function in the thoracic muscles only (*Act88F >CapaR^KO^*) failed to: (i) reduce locomotor activity (flies with rescued Capa signaling in muscles, however, appear to be hyperactive), (ii) disrupt systemic energy homeostasis apart from reducing glycogen levels (rescue animals appear hyperglycemic); (iii) impair food intake; or (iv) induce significant mortality compared to the controls (Fig. S5a-h). Consequently, impairing Capa/CapaR signaling exclusively in flight muscles does not recapitulate the full extent of phenotypes of global muscle knockout, suggesting that direct Capa-induced changes in skeletal muscle performance is unlikely to be the major cause to the acute mortality observed in *how^ts^>CapaR^KO^* flies.

The skeletal musculature imposes large energy demands on the organism due to its large mass and high metabolic rate. We therefore asked whether reduced systemic energy levels cause the observed decline in locomotor activity. First, we assessed the tissue-specific distribution of glycogen (the main energy storage form in muscles) as a functional readout of skeletal muscles’ energy status using the periodic acid-Schiff (PAS) staining method^35^. Remarkably, PAS staining was almost completely abolished in the muscles and to a lesser extent in the fat body (but not in brain) of muscle-specific *CapaR* knockout flies (*how^ts^>CapaR^KO^*). In addition, this effect was partially rescued by co-expressing wild-type CapaR (Fig. 6e). These data confirmed that the major glycogen storing tissues (i.e. the fat body and skeletal muscles) are depleted of glycogen, which together with the generally lean phenotype of *how^ts^>CapaR^KO^* animals (Fig. 5i-l), implies that they are unable to meet the high energy demands of contracting muscles. In line with this hypothesis, we observed that the genes encoding the rate-limiting glycogenolytic enzymes Glycogen phosphorylase (*GlyP*) and Debranching enzyme (*AGL*) are upregulated in *CapaR* knockout muscles, while the glycogenic enzymes glycogen synthase (*GlyS*) and Branching enzyme (*AGBE*) are significantly downregulated (Fig. 6f), indicating that the skeletal muscles are adapting metabolically by increasing glycogen mobilization in an attempt to meet autonomous energy demands.

Given that muscle performance depends critically on effective calcium cycling mechanisms, and that Ca^2+^-pump activity is highly sensitive to cellular energy levels^36^, we rationalized that energy depletion in muscle cells may compromise Ca^2+^ dynamics. To test this hypothesis, we quantified cytosolic Ca^2+^ levels during steady-state and Capa-1 stimulation in muscle-specific *CapaR* knockdown animals with reduced muscle performance. Strikingly, these data showed that not only does *CapaR* depletion abolish the Capa-induced increase in intracellular Ca^2+^ levels, but the unstimulated basal [Ca^2+^]_i_ levels were almost 50% increased relative to the control (Fig. 6g-h). This observation suggests that the transport machinery responsible for reestablishing resting Ca^2+^ levels is compromised. We therefore investigated a potential role of the *Drosophila* plasma membrane calcium ATPase (*PMCA*) in muscle function – an efflux transporter crucial to maintenance of resting [Ca^2+^]_i_ levels and for regulating the excitation-contraction coupling process in muscles^37^ – and found that targeted knockdown of *PMCA* in muscles phenocopied the dysregulated [Ca^2+^]_i_ levels, impaired locomotor activity and mortality phenotypes observed in *how>CapaR^KO^* flies (Fig. S5i-l). Consistent with these observations, transcript levels of *PMCA* were significantly reduced in *CapaR* knockout flies compared to control and rescue animals (Fig. 6i), suggesting that Ca^2+^ dysregulation in *how>CapaR^KO^* skeletal muscles is at least partly linked with reduced PMCA activity. Overall, our data suggest that the progressive loss of muscle function observed in Capa/CapaR impaired animals is unlikely to be caused by direct Capa actions on skeletal muscles. Rather, our data point to a model in which depletion of internal energy stores and reduced Ca^2+^-pump activity disrupts Ca^2+^-dynamics in muscles, creating a muscular dystrophy-like phenotype analogous to that observed in human myopathies^38^.

### Nutrients and water induce release of Capa peptides

Our data imply that Capa/CapaR signaling plays a central role in coordinating postprandial homeostasis, such as by increasing gut peristalsis, facilitating nutrient absorption and increasing renal secretion. We therefore tested if the *Capa*-expressing neurons are sensitive to internal signals related to water and nutrient availability. This hypothesis is consistent with the observation that Capa peptides accumulate in Va neurons during desiccation stress, but are subsequently released during recovery^39^. We thus exposed flies to either desiccation or starvation for 24 hours followed by transfer to media with different water and nutrient compositions and applied complementary approaches to measure neurosecretory activity (Fig. 7a). First, we quantified *Capa* transcript levels as well as intracellular Capa prohormone levels in Capa^+^ neurons. Secondly, we used an *in vivo* calcium reporter (CaLexA) that translates sustained neural activity into nuclear import of the chimeric transcription factor LexA::VP16::NFAT and thus to subsequent LexAop-GFP expression^40^. Consistent with previous findings, these data showed that *Capa* mRNA levels and immunoreactivity were significantly increased in Va neurons, but not SEG neurons, of desiccated flies compared to control, yet returned to steady-state levels or lower 4 hours after refeeding or rehydration (Fig. 7b-c; Fig. S6). As expected, we observed inverse changes in activity in flies carrying *Trp-GAL4* and *UAS-mLexA::VP16::NFAT* in Capa-expressing neurons (Fig. 7d). These data suggest that the Va neurons are inactive during periods of water and/or nutrient restriction but are induced to release Capa neuropeptides at high rates following both refeeding and rehydration. Interestingly, however, long-term hydration of rehydrated flies led to gradual accumulation of Capa peptides, and a concomitant reduction in GFP intensity over the ensuing 24 hours (Fig. 7c-d), indicating that water ingestion alone is insufficient to maintain stimulation of Capa-Va neuronal activity in hydrated flies.

**Figure 7.**
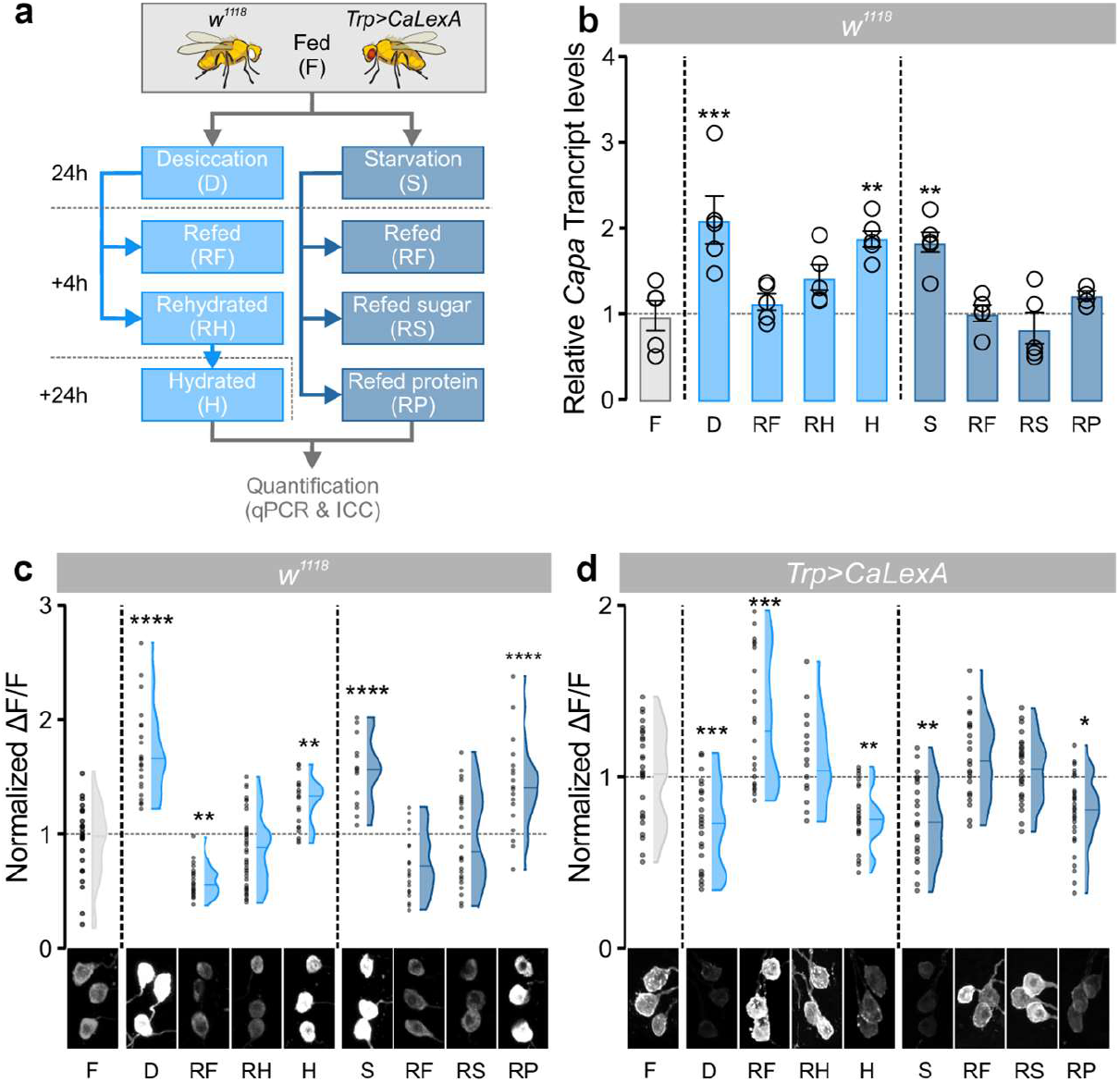
Environmental cues modulating Capa^+^ Va neuron activity. **a.** Experimental design for the environmental exposures of wild-type (*w^1118^*) and *Trp>CaLexA* flies. **b.** Transcript levels of *Capa* gene expression relative to *RpL3* in Va neurons (thorax samples) of flies exposed to the environmental conditions indicated (one-way ANOVA; n=5). **c.** Violin plots and raw data points of immunofluorescence quantifications of intracellular Capa precursor levels (n=18-52) and **d.** CaLexA-induced GFP expression (n=18-36) in Capa^+^ Va neurons with representative images from all conditions (one-way ANOVA).

Since starvation affects Capa release, we tested whether the Capa^+^ neurons were sensitive to nutritional cues. In flies exposed to nutrient-deprivation for 24 hours, we observed a significant increase in *Capa* mRNA levels and Capa immunoreactivity and a concomitant reduction in GFP reporter expression in the Va neurosecretory cells, with both Capa and GFP intensities returning to steady-state levels following refeeding on standard medium (Fig. 7b-d). These data demonstrate that the Capa^+^ Va neurons, and again not the SEG neurons (Fig. S6), are responsive to signals related to internal nutrient availability. To further identify the dietary factors that regulate Capa release, we transferred starved flies to media that contained either sucrose and yeast as exclusive sources of carbohydrates or amino acids, respectively. Interestingly, refeeding sugar, but not amino acids, reverted anti-Capa and GFP fluorescence levels similar to the ones observed in fully fed conditions (Fig. 7c-d), suggesting that the Va neurons are specifically activated in response to dietary sugars or their metabolites. Together, our data show that the different populations of Capa^+^ neurons are differentially activated: only the three pairs of Capa^+^ Va neurons relay information about the internal availability of water and sugar levels, because only these neurons release Capa peptides into the circulation in response to environmental conditions known to affect the internal abundance of these nutrients.

### Capa-mediated regulation of AKH secretion controls systemic metabolic homeostasis

The finding that *Capa* neurons respond directly to internal sugar levels implies that Capa peptides relay nutritional information to exert metabolic control. This prompted us to explore a potential interaction between Capa signaling and hormonal systems controlling systemic energy balance. As in mammals, the *Drosophila* insulin-like peptides (DILPs) and AKH (a glucagon analogue) act as the key counter-regulatory hormones in controlling carbohydrate metabolism^8,30,41^. Although we were unable to detect GFP expression in the adult insulin-producing cells (Fig. 2a), we surprisingly found reporter-gene expression (*CapaR>GFP*) in the AKH-producing neurosecretory cells in the CC; the sole site of AKH synthesis and release^30^. Furthermore, we observed specific and displaceable Capa-1-F binding overlapping with APC-targeted GFP expression (*Akh>GFP*) (Fig. 8a-b). These data suggest that Capa peptides regulate energy metabolism by directly modulating the AKH signaling pathway. Because the primary function of AKH is to induce lipid catabolism, which affects organismal resistance to starvation, we measured survival under starvation in animals with reduced *CapaR* expression in the APCs to test whether Capa regulates AKH secretion. Flies with adult-specific knockout of *CapaR* in the APCs (*Akh^ts^>CapaR^KO^*) showed a marked hypersensitivity to nutrient deprivation compared to age-matched controls (*Akh^ts^/+*), an effect previously associated with increased AKH release^30^; we furthermore observed that this effect was fully rescued in flies additionally carrying a *UAS-CapaR* transgene (Fig. 8c). These data show that Capa mediates adult-specific nutritional regulation of AKH signaling, which is necessary to sustain organismal survival during nutrient deprivation.

**Figure 8.**
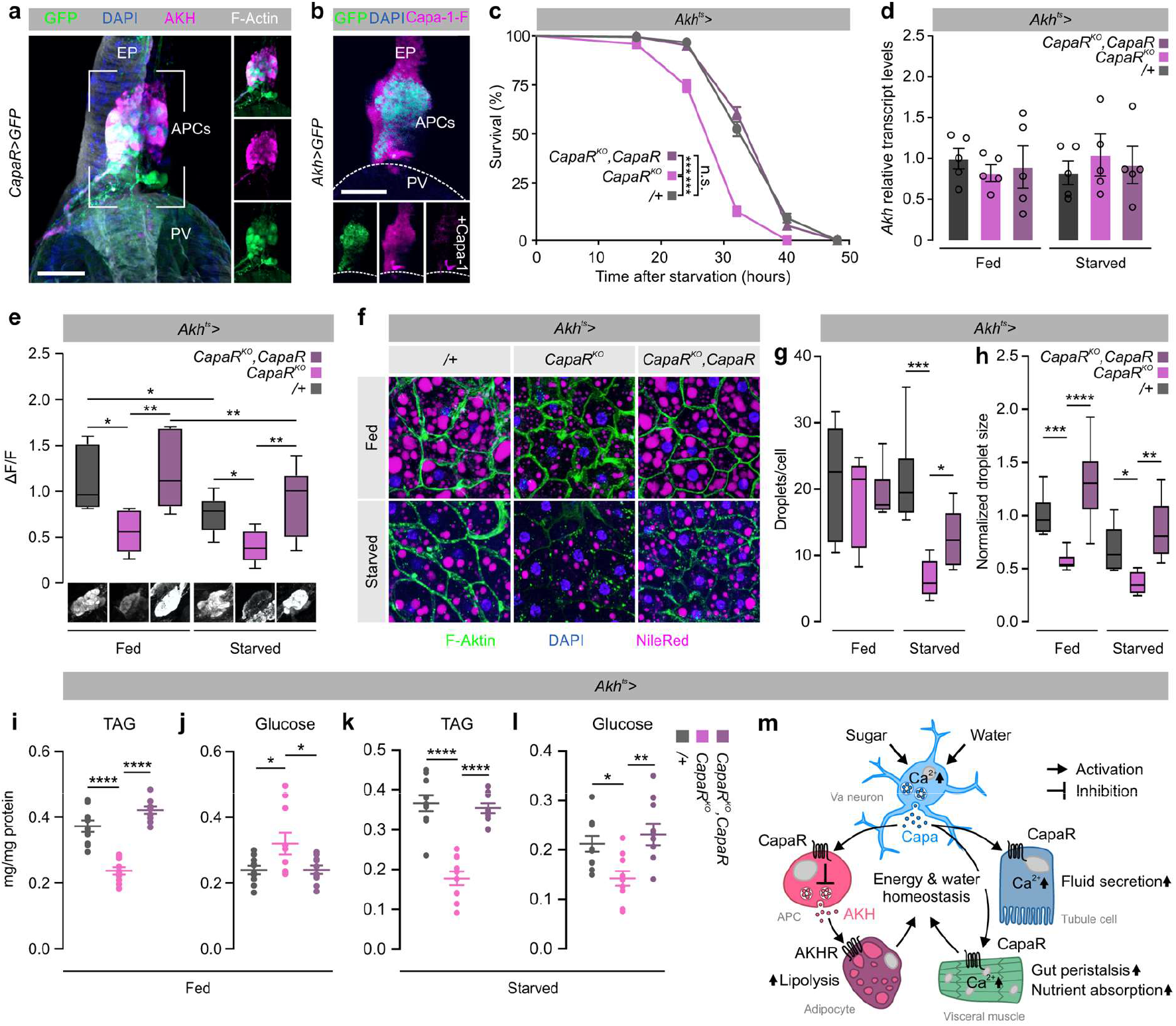
Systemic Capa signaling regulates metabolic homeostasis via AKH activity. **a.** AKH immunofluorescence (magenta) colocalizes with *CapaR>* driven *mCD8::GFP* expression (green) confirming CapaR localization to the APCs. Scale bar 10 μm. **b.** Application of Capa1-F (magenta) shows specific and displaceable binding to *Akh>GFP* (green) positive cells in the APCs. Scale bar 10 μm. EP, esophagus; PV, proventriculus; APCs, AKH-producing neurosecretory cells. **c.** *CapaR* knockout in the APCs (*Akh>CapaR^KO^;* n=208,) results in significantly reduced resistance to starvation compared to parental controls (+/; n=213), while co-overexpression of *CapaR* (*Akh>CapaR^KO^;CapaR;* n=247,) rescues the survival phenotype (log-rank test). **d.** Transcript levels of *Akh* relative to *RpL32* in the APCs from the different genetic backgrounds (one-way ANOVA; n=5). **e.** Immunofluorescence measurements of intracellular AKH levels in the APCs from animals exposed to either *ad libitum* feeding (fed) or caloric deprivation (starved) for 24 hours, probed with an anti-AKH antibody (one-way ANOVA; n=6-15). **f.** Lipophilic dye Nile red staining of adult dissected fat bodies from fed and starved control (/+), knockout (*CapaR^KO^*) and rescue flies (*CapaR^KO^, CapaR*). **g.** Quantification of lipid droplets/cell and **h.** lipid droplet size in fed and starved flies with the different genetic backgrounds (one-way ANOVA; n=6-8). **i.-l.** Quantification of whole-body TAG and **j.** glucose in fed and starved animals of the different genotypes (one-way ANOVA; n=10). **m.** Working model for the systemic actions of Capa/CapaR signaling in mediating water and osmotic homeostasis in adult *Drosophila.* Capa^+^ Va neurons respond to internal sugar and osmotic cues by activating two separate pathways: one stimulating intestinal and renal activities to promote gut contractions, nutrient absorption and fluid secretion, and another to restrict glucagon-like AKH release from the APCs and subsequent lipolytic activity in adipose tissue.

To further probe the interactions between AKH and Capa signaling, we compared *Akh* transcript and intracellular AKH protein levels upon APC-specific *CapaR* knockout in flies exposed to either *ad libitum* feeding or caloric deprivation. Neither starvation nor *CapaR* manipulation altered *Akh* expression (Fig. 8d). However, starvation induced a marked decrease in intracellular AKH levels in the APCs in all genotypes, consistent with the known induction of AKH release by starvation^30^ (Fig. 8e). Moreover, in both fed and starved conditions we also observed that *CapaR* knockout (*Akh^ts^>CapaR^KO^*) consistently lowered intracellular AKH protein levels relative to control, which was reverted to baseline levels in rescue animals (*Akh^ts^>CapaR^KO^, CapaR*) (Fig. 8e). These data demonstrate that CapaR activation in the APCs act to repress release of AKH into circulation in a nutrient-dependent manner, and that the Capa neuroendocrine pathway thus exerts metabolic control by modulating AKH activity. Such a function is furthermore consistent with the sugar-induced release of Capa from the Va neurons given that AKH release should be inhibited during sugar-replete states, in part via Capa/CapaR, in order to prevent lipolysis.

Given that the main downstream action of AKH is to regulate lipolysis in adipose tissue, we investigated the metabolic status of *CapaR*-deficient animals by assessing their lipid reserves in the adult fat body. Using the lipophilic dye Nile Red to direct visualize lipid contents^42^, we observed a significant but rescuable reduction in fat body lipid-droplet size in *CapaR*-knockout animals under both fed and starved conditions; starvation also induced a partially rescuable decrease in lipid-droplet number in *Akh^ts^>CapaR^KO^* animals (Fig. 8f-h). Consistent with these observations, we detected a significant reduction in organismal energy reserves in both fed and starved animals compared to control, as indicated by decreased whole-body TAG levels in APC-specific *CapaR* knockout animals (Fig. 8i,k). In addition, fed flies lacking *CapaR* in the APCs displayed a pronounced hyperglycemia relative to control and rescue animals, consistent with hyperactivation of AKH signaling in these flies. In contrast, starved flies exhibited a decrease in glycemic levels, likely due to an accelerated depletion of organismal energy reserves (Fig. 8j,l). Altogether, our results identifies an intertissue program involving Capa^+^ Va neurons, which sense internal sugar and water availability and signal to neurosecretory cells in the CC to control AKH-mediated catabolic processes in adipose tissue, adding another layer of regulation to the complex control of energy homeostasis in the adult fly.

## DISCUSSION

### Between what goes in and what goes out: Capa signaling directs post-ingestive intestinal physiology and excretory behavior

Our work has identified a system in which Capa peptides released by six Va neuroendocrine cells of the ventral nerve cord activate their receptor CapaR localized to visceral muscles to control intestinal motility and food transit. Activating the Capa circuit increases intestinal transit causing flies to excrete large amounts of dilute deposits, whereas inactivation of CapaR in gut muscle cells results in reduced gut motility and excretion. These physiological effects are linked with the strategic localization of CapaR expression to visceral muscles in distinct regions of the gut that contain valves and sphincters^22^, implying that Capa peptides are part of a digestive program by which the transit of intestinal contents is sensed and modulated. Consistent with this idea, genetic inhibition of gut peristalsis and intestinal passage appear to increase the amount of fluid absorbed by the gut, which leads to a concentration of waste products and causes a constipation-like phenotype analogues to the effects observed in humans suffering from gastrointestinal motility disorders^25^. This manipulation also frequently associates with intestinal distention, which incidentally helps explain the reduced food intake by these flies, given that enteric neurons convey stretch-dependent feeding-inhibitory signals^43,44^. Our work and previous studies^12,45^ are therefore consistent with a model in which the synchronized control of renal secretion and gut motility by the Capa circuit helps synergize the homeostatic control of water balance and waste excretion, which shows striking functional homology to the actions of NmU signaling in humans^13^. Intriguingly, the coupled control of renal and gut functions appears to have been co-opted during evolution since other hormones that exert systemic control of diuresis – such as kinin, Diuretic Hormone 31 (DH31) and Diuretic Hormone 44 (DH44) – have also been reported to modulate feeding behavior and intestinal contractions^6,44,46,47^. The convergence of multiple neuropeptide signaling pathways on organ functions critical to controlling postingestive physiology, suggests a fundamental need for coupling food intake with fluid and waste excretion. How these endocrine networks interact to regulate these vital processes remains to be examined.

In addition to compromising intestinal fluid balance and excretion, we also found that silencing Capa-mediated visceral contractions disrupt the integrity of the midgut acidic zone. The acidic zone consists of acid-secreting copper cells that are functionally related to the gastric parietal cells of the mammalian stomach, and this region is necessary to facilitate the digestion and absorption of macronutrients and metals^26,48,49^. Although we cannot exclude that the loss of acidity is due to reduced copper-cell acid secretion, our data suggests that CapaR expressing visceral muscles at the anterior and posterior junctions of this region serve a sphincter-like role that is necessary to maintain gut compartmentalization and optimal pH of the CCR; a role strikingly similar to that played by the esophageal and pyloric sphincters in humans. This idea is consistent with the morphological constrictions of the gut tube at either end of the CCR and is supported by a recent study demonstrating that ablation of enteroendocrine derived DH31, which controls food passage into the acidic zone in *Drosophila* larvae, leads to inappropriate mixing of acidified and non-acidified gut contents^48,50^. Our data thus point to a mechanism in which nutrient-dependent Capa release helps maintain functional gut compartmentalization in order to promote and optimize nutrient digestion and uptake following food consumption.

### Capa-mediated control of systemic metabolism affects locomotor activity and survival

Silencing of Capa/CapaR signaling in muscles induces a complex metabolic phenotype characterized by lipodystrophy and hypoglycemia accompanied by hypophagia, locomotor deficits, and reduced organismal lifespan. How can we reconcile these phenotypes? Several lines of evidence indicate that impaired nutrient uptake is a conserved mechanism reducing organismal longevity^51,52^, raising the possibility that these phenotypes arise as a consequence of malnutrition and poor intestinal absorption. This idea is consistent with the gradual decrease in whole-body energy stores following conditional *CapaR* knockout in visceral muscles and is further supported by the observation that survival can be enhanced by exposing these flies to higher nutrient concentrations. Concordantly, our data show that the loss of internal metabolic stores correlates with the progressive decline in physical activity. These results strongly suggest that the age-dependent decline in locomotor activity derives from impaired energy homeostasis affecting the musculature. Intriguingly, locomotor deficits are known to impair foraging behavior and food intake^53^, suggesting that the reduced muscle performance observed in Capa/CapaR deficient animals may contribute to the hypophagia exhibited by these animals, causing further malnutrition and ultimately death. Altogether, our data support a model in which nutrient absorption is one of the key aspects of Capa-mediated control of adult metabolic homeostasis – a function that is essential to sustain physical activity, feeding and survival,

### Coupled sensing of internal sugar and water abundance for maintaining homeostasis

Our study suggests that the Capa^+^ Va neurons, but not the SEG neurons, secrete Capa peptides in the presence of high internal water and sugar levels, while desiccation and caloric deprivation reduce release and thus results in Capa hormone accumulation. These data connect homeostatic variations in internal water and sugar availability to Capa signaling, and remarkably show that the Capa Va and SEG neurons are activated by different physiological signals. Yet, how might this type of regulation be adaptive? One possibility is that food and water consumption change the internal metabolic status of animals (in a manner dependent on food composition), requiring appropriate activation of specialized organs to restore homeostasis. For example, feeding in *Drosophila* is associated with fluid uptake, which needs to be balanced by renal excretion to avoid acute effects; prolonged inactivation of diuresis can cause severe fluid retention, resulting in cuticle rupture and death^22^. Conversely, nutrient and water deprivation induce osmotic perturbations requiring compensatory actions to reduce organismal water loss; inhibiting tubule and intestinal activities promotes fluid retention and increase survival under desiccating conditions^45,54^. Our study thus provides evidence of a mechanism by which homeostatic deviations in internal osmolality may be reported by the Capa^+^ Va neurons, which provide systemic feedback regulation on intestinal and renal activities to restore fluid and ion balance.

Our finding that Capa^+^ Va neurons are potently activated by high hemolymph sugar levels – a conserved signal of energy surfeit – and that Capa modulates AKH release and thus govern lipolytic activity in the fat body, surprisingly implicate Capa action in glycemic control. Our results consistently show that impaired Capa signaling in the APCs due to *CapaR* elimination induces a marked hypersensitivity to starvation; a phenotype that is also associated with a significant increase in AKH secretion, reduced fat body energy reserves, and hyperglycemia (Fig. 8c-j). Whereas the impact of insulin regulation on carbohydrate metabolism is well studied in both humans and flies, less is known about AKH/glucagon. Our study reveal a neuroendocrine mechanism in which Capa peptides released from the CNS exert metabolic control by providing feedback regulation on AKH-mediated lipolysis in the fat body to stabilize circulating sugar levels. Intriguingly, similar suppression of a metabolic hormone was recently reported for NmU signaling in mammals^55^, implying that vertebrates and invertebrates may share evolutionarily conserved mechanisms for regulating sugar metabolism.

A key requirement of homeostatic regulation is the ability to sense deviations in internal abundances and to initiate compensatory actions to restore balance. Although we do not know whether the Capa^+^ Va neurons are directly sugar- and osmosensitive, or alternatively if such information is encoded by other neurons, it is surprising that cues related to both sugar and water availability converge on the same group of neurons. Strikingly, a recent study found that expression of AKHR, and the TRPV channel, Nanchung, by the same group of interoceptive neurons underlie their ability to direct monitor internal signals of nutrient and water availability^7^. Similarly, sugar and water reward have also been found to be processed by several other subsets of neurons in the brain, in a manner independent of gustatory sensory activation^6,8,56,57^. These data imply that several internal nutrient sensors in the *Drosophila* brain communicate homeostatic needs for sugar and water independently, and that Capa signaling may interact with other signaling networks to couple consumption with the regulation of absorptive and excretory organs to restore balance. Indeed, it is likely that in the face of complex environmental challenges, multiple mechanisms converge to ensure a robust organismal response to diverse stressful conditions to sustain animal survival.

In sum, our work has uncovered a key role for Capa signaling in coordinating organ-specific post-prandial responses that are essential to sustaining adult fly survival. This interorgan signaling system consists of a CNS/gut/renal/fat body signaling module, which operates via a discrete population of sugar- and osmosensitive Capa^+^ neurons that release Capa peptides into circulation to control gut peristalsis, nutrient uptake, renal secretion and fat body lipolysis, to ultimately promote internal osmotic and metabolic homeostasis following sugar and water ingestion (Fig. 8m). Overall, the striking functional conservation observed between insect Capa and mammalian NmU signaling further emphasizes the unique power of the *Drosophila* model system as a tool that may provide valuable insights into the complex regulation of metabolic and energy homeostasis in humans.

## MATERIALS AND METHODS

### Fly stocks

*UAS- CapaR^RNAi^ (2727), UAS-mCD8::GFP* (5137), *UAS-mLexA-VP16-NFAT* (66542), *UAS-GCaMP6s* (42749), *elav-GAL4* (458), *Cg-GAL4* (7011), *201Y-GAL4* (4440), *c564-GAL4* (6982), *how(24B)-GAL4* (1767), *Akh-GAL4* (25684), *UAS-Tubulin-Gal80ts* (7019), *Act88F-GAL4* (38461) and *Trp-GAL4* (49296) flies were obtained from Bloomington *Drosophila* Stock Center (Indiana University, Bloomington, USA). Moreover, *UAS-CapaR^RNAi^ (105556*) and *PMCA^RNAi^* (101743) transgenic fly lines were obtained from the VDRC RNAi stocks (Vienna *Drosophila* Resource Center, Austria). *Tinman-GAL4* was kind gift from Dr. Manfred Frasch (University of Erlangen-Nuremberg, Germany) and *Hand-GAL4* was kind gift from Dr. Achim Paululat (University of Osnabrück, Germany). The *CapaR-GAL4, CapaR-GAL4; UAS-GFP::aequorin, how(24B)-GAL4; UAS-GFP::aequorin*, and *Uro-GAL4* drivers were previously generated in-house. In addition, a viable doubly homozygous *CapaR* RNAi (II, III) fly line was generated using standard genetic techniques. All fly lines were maintained on a standard *Drosophila* cornmeal medium at 25°C, 50-60% humidity under a 12-hour:12-hour light-dark photoperiod. Standard fly crosses were performed at 25°C. Flies carrying the *TubulinGal80^ts^* transgene encoding a temperature-sensitive transcriptional repressor were raised at 18°C and subsequently sorted for males and females upon eclosion. After sorting, flies were kept for another 24 hours at 18°C before being switched to 30°C for maturation. Only male flies were used for experimentation. *w^1118^* was used as wild-type control.

### *Drosophila* transgenesis

The ORF of *CapaR* (*CG14575*) was amplified from whole *w^1118^*fly cDNA as template and cloned into pUAST attB vector using *NotI* and *KpnI* whose recognition sites are included on the primers (Table S1), then integrated on the second chromosome by site-directed insertion using the phiC31 integrase and an attP landing site carrying recipient line, *y[1] w[1118]; PBac{y[+]-attP-9A^58^ VK00018}* (Bloomington *Drosophila* Stock Center #9736). The tissue-specific *CapaR* CRISPR^KO^ construct was cloned into pCFD6 vector (www.crisprflydesign.org, Addgene #73915) according to the website’s instruction. In this vector, we cloned four independent gRNA constructs designed to target non-coding regions (introns and 3’UTR; see primers listed in Table S1) of the *CapaR* genomic DNA sequence. After sequencing, the construct was inserted on the third chromosome in an *attP2* landing site carrying recipient line, *y,w*, *P(y[+].nos-int. NLS); P(CaryP)attP2* (gift from Dr. Diogo Manoel, Instituto Gulbenkian de Ciência, Portugal). Germline transformation was carried out in-house.

### Survival assay

Flies from each genetic background were collected within 24 hours after eclosion and maintained on standard *Drosophila* cornmeal medium (app. 30 flies per vial) at either 18, 25 or 30°C at 50-60% humidity with a 12-hour:12-hour light-dark photoperiod. In all conditions, survivors were transferred into new vials every three days. Each vial was then observed for dead flies (no movement after gently tapping the vial) every 24 hours for 10 days. Data was expressed as per cent survival over time with 213-793 flies counted for each genotype and condition.

### Quantitative real-time PCR analysis

Total RNA was extracted from whole animals or dissected tissues using the RNeasy Mini Kit (QIAGEN) with DNase treatment to remove contaminating genomic DNA. RNA was quantified using a NanoDrop spectrophotometer (Thermo Scientific) and reverse transcribed into cDNA using iScript Reverse Transcription Supermix (Bio-Rad). Quantitative real-time PCR (qPCR) was performed using the QuantiTect SYBR Green PCR Kit (Qiagen) and the Mx3005P qPCR System (Agilent Technologies) with transcript levels normalized to *RpL32* or *RpL3* reference genes and expressed as fold change compared to controls ± s.e.m (n=5). The primers used are listed in Table S2.

### Whole fly paraffin-embedded sections and immunostaining

Paraffin sections were made according to a modified protocol previously described^59^. In brief, *CapaR>GFP* flies were fixed in 4% (wt/vol) paraformaldehyde (PFA) for 30 minutes, and then dehydrated through a graded series of ethanol and xylene, before being embedded in paraffin. Next, ~8 μm thick paraffin sections were cut on a Leica RM 2235 microtome (Leica Biosystems, Nussloch, Germany), and the sections were transferred to individual glass microscope slides for subsequent use. For immunostainings, the sections were deparaffinized in Histo-Clear (National Diagnostics, USA) for 15 minutes, rehydrated through a graded series of alcohol (99%, 90% and 70% for 10 minutes each step), and rinsed in PBS. The sections were then incubated in blocking buffer (PBS with 0.1% Triton X-100 and 2% normal goat serum), containing polyclonal Alexa Fluor 488-conjugated rabbit anti-GFP (1:200; ThermoFisher, #A21311) and DAPI (4’,6-diamidino-2-phenylindole, 1 μg ml^-1^; ThermoFisher, #D1306) overnight at 4°C. Finally, the sections were washed repeatedly in PBS and mounted in Vectashield (Vector Laboratories Inc, CA, USA). Image acquisition was performed on an inverted Zeiss LSM 880 confocal laser-scanning microscope (Zeiss, Oberkochen, Germany) and processed with CorelDraw X8.

### Immunocytochemistry

Immunocytochemistry was performed as previously described^60^. In brief, *Drosophila* tissues were dissected in Schneider’s insect medium (Invitrogen, CA, USA), fixed in 4% (wt/vol) paraformaldehyde in PBS for 30 min at room temperature and washed twice for 1 hour in blocking buffer. Tissues were then incubated in primary antibodies in blocking buffer overnight, followed by overnight incubation in secondary antibodies in blocking buffer at 4°C. Counter staining with DAPI and/or Rhodamine-coupled phalloidin (1:100; ThermoFisher, #R415) was performed where appropriate. The primary antibodies used were Alexa 488 conjugated goat anti-GFP (1:500), rabbit anti-CapaR^12^ (1:500), rabbit anti-Capa precursor peptide^10^ (1:500) and rabbit anti-AKH (raised against the secreted portion of the peptide^30^), a generous gift from Dr. Jae Park, University of Tennessee, US (1:500). The primary antibodies were visualized with goat anti-rabbit Alexa Fluor 488, 555 or 594 (1:500; ThermoFischer #A32731, #A21429 or #R37117). For staining of intracellular lipid droplets in the fat body, adult adipose tissue was fixed as described above and the lipophilic dye Nile red (2.5μg/ml) was applied for 30 minutes after which tissues were rinsed with PBS. All samples were mounted on poly-L-lysine coated 35 mm glass bottom dishes (MatTek Corporation, MA, USA) in Vectashield and imaged as previously described. Retained intracellular Capa prohormone and AKH peptide levels as well as lipid droplet size and number were calculated using the FIJI software package.

### Peptide synthesis and *ex vivo* receptor-binding assay

*Drosophila* Capa-1 and Capa-2 (both with and without an N-terminal cysteine) were synthesized by Cambridge Peptides (Birmingham, UK), and subsequently coupled to high quantum yield fluorophores via a cysteine-linker to make fluorescent TMR-C_5_-maleimide-GANMGLYAFPRVamide (Capa-1-F), and Alexa488-C_5_-maleimide-ASGLVAFPRVamide (Capa-2-F). Fluorescent Capa peptides were applied to tissues in an *ex vivo* receptor-binding assay as previously described^16^. In brief, tissues of interest were dissected from cold anesthetized animals and mounted on poly-L-lysine coated glass bottom dishes before being setup in a matched-pair protocol. One batch was incubated in 1:1 (vol/vol) mix of *Drosophila* saline and Schneider’s medium (artificial hemolymph; AHL) containing DAPI, while to the other batch was additionally added 10^−7^ M of Capa-1-F. The batch without the labeled peptide was used to adjust baseline filter and exposure settings to minimize background during image acquisition on a Zeiss LSM 880 confocal microscope. Specificity of binding was additionally verified by competitive displacement of the labeled ligand with unlabeled peptide at a concentration of 10^−5^ M.

### Real-time measurements of intracellular calcium release

Cytosolic calcium measurements were performed according that previously described^61^. Tissues were dissected from flies expressing *UAS-GFP::aequorin* driven by *CapaR-GAL4* or *how-GAL4.* For each sample, 20-30 live intact adult tubules, proventriculus, midgut, hindgut, brain, gonads, salivary gland, legs or thoraxes were transferred to 5 ml Röhren tubes (Sarstedt AG & Co., Nümbrecht, Germany) in 175 μl Schneider’s medium and subsequently incubated in the dark with 2 μl coelenterazine in ethanol (final concentration of 2.5 μm) for 3 hours to reconstitute active aequorin. Real-time luminescence was measured on a Berthold Lumat LB 9507 luminometer (Berthold technologies, Bad Wildbad, Germany). A stable baseline was established prior to both mock injection and subsequent injection with Capa-1 peptide (final concentration 10^−7^ M), and the luminescence was measured in the ensuing period. After each experiment, undischarged aequorin was measured by permeabilizing the cells with 300 μl lysis buffer (1% Triton X-100 and 0.1 M CaCl_2_). Real-time calcium concentrations ([Ca^2+^]_i_) throughout the experiments were then back-calculated with an in-house Perl routine.

### Gut motility assay

Guts from control and *CapaR* knockdown flies were dissected in artificial hemolymph (AHL) with minimal disruption of attached tissues and without removing the head. Individual exposed guts were next pinned onto a Sylgard-lined petri dish, with fine tungsten pins through the proboscis and a small piece of cuticle attached to the end of each gut and bathed in 100 μl of AHL. Following a 5-minutes acclimation period, each gut was then recorded with a Leica IC80 HD camera mounted on a Leica M50 stereomicroscope for 3-minutes, before the solution was exchanged with 100 μl of AHL and recorded for another 3-minutes. This was done to correct for potential artefacts on gut contraction frequency associated with the exchange of solutions. Next, the AHL solution was replaced with 100 μl of 10^−7^ M of either Capa-1 or Capa-2 peptide in AHL, and tissues recorded for an additional 2 x 3-minutes. Contractions of the entire gut were visually counted post image acquisition, and the number of contractions was normalized to the number of contractions observed in the initial AHL solution with n= 6-8 guts counted for each genotype.

### Calcium imaging

To measure muscle activity upon Capa peptide stimulation, guts from *CapaR>UAS-GCaMP6s* flies were dissected in AHL and immobilized by mounting in poly-L-lysine coated dishes. Each gut was recorded for 500 frames in total (512 x 512 pixels; one frame per 5 seconds); the initial 100 frames were recorded prior to Capa peptide stimulation, the next 200 frames were recorded during stimulation with 10^−7^ M Capa-1, and the final 200 frames were recorded during wash-out.

### Quantitative analysis of defecation behavior

Analysis of defecation behavior was performed according to a modified protocol^22^. Standard *Drosophila* food was prepared and allowed to cool to roughly 65°C before it was supplemented with 0.5% (wt/vol) Bromophenol blue (BPB) sodium salt (B5525, Sigma). Both gut transit and excreta quantification experiments were performed on batches of 10 flies, which were placed in standard fly vials or 50-mm Petri dishes containing BPB-dyed food. For gut transit measurements, the flies were transferred to fresh vials every hour and the number of excreta manually scored. For Petri dishes, digital images of Petri lids were obtained using an photo scanner (Konica C454e), which were processed in ImageJ before being analyzed (including depsit number, size, and dye intensity) using the TURD software package^21^. Dissections of dye-containing intestines were performed in AHL, leaving the head and posterior cuticle intact to prevent leakage of gut contents.

### Scanning electron microscopy and quantitative elemental microanalysis

Deposits were collected and transferred to individual aluminium stubs. Samples were then examined in a Zeiss Sigma variable pressure analytical scanning electron microscope (Carl Zeiss, Oberkochen, Germany) in combination with AZtecEnergy microanalysis software (Oxford instruments, Oxford, UK). Element microanalysis and acquisition of composition maps was carried out using high accelerating voltages (≥20kV) in combination with electron backscatter (BSE) and both energy-dispersive X-ray spectroscopy (EDS) and wavelength-dispersive X-ray spectroscopy (WDS), providing both structural and quantitative measures for excretion composition at high resolution.

### Feeding assays

Groups of flies were starved for 4 hours, and then transferred to normal food containing 0.5% BPB for another 4 hours, followed by freezing the flies at −80 °C. Groups of five flies were transferred to 2 mL tubes and were homogenized in 100μl PBS + 0.1% Triton X-100 using a TissueLyser LT (Qiagen). The samples were then spun down at 12,000 rpm and the supernatant transferred to fresh vials and the absorbance of each sample at 603 nm was measured using a NanoDrop. The resulting absorbance values were then subtracted the mean background value obtained from unfed flies (n=10), and the data were normalized to control. A total of n=22-27 samples were measured for each genotype.

For analysing feeding behavior in individual flies, animals were starved for 4 hours, before being placed in a *Drosophila* breeding chamber separated by a divider. Following a 5 min acclimation period, the divider was removed allowing the fly access to small drop of 5% (wt/vol) glucose in in ddH_2_O. The chambers were filmed in the ensuing 10 minute period, and the time spent in contact with food during this period was quantified post acquisition with n=10-12 for each genotype.

### Nutrient rescue

An artificial diet chiefly consisting of a yeast-sugar medium was used for a nutritional rescue assay. The medium consisted of 80 g/l yeast and 90 g/l sucrose boiled with 1% agar, 4.8% propionic acid and 1.6% methyl 4-hydroxybenzoate (all components from Sigma). Media with lower nutrient concentrations were made by keeping sucrose concentration constant but reducing yeast concentrations to 16 g/l and 0 g/l, respectively. Up to 30 animals were transferred into vials containing the appropriate medium, and fly mortality was counted every 12 hours for the ensuing 10 days with a total of n=251-323 flies used per condition.

### Nutrient-level assays

Four days after transfer to the restrictive temperature (30 °C), flies were collected and frozen at −80 °C in groups of ten animals, and subsequently homogenized in 300 μl PBS + 0.5% Tween-20 (PBST) using a TissueLyser LT (Qiagen). Next, 2 μl was used to quantify protein content using the Bradford assay for subsequent normalization of nutrient levels. The samples were then heat-inactivated at 70 °C for 5 minutes, centrifuged to pellet debris for 3 seconds at full speed, before the supernatant was transferred to fresh vials. For triacylglyceride measurements, 2 μl of the supernatant was incubated with 4 μl Triglyceride Reagent (Sigma, T2449) and 6 μl PBST at 37 °C for 30 minutes; next, 20 Free Glycerol Reagent (Sigma, F6428) was added, and the reaction was incubated at 37 °C until color development occured (usually 30-60 min). Absorbance at 540 nm was then measured. For glycogen measurements, 1 μl homogenate was mixed with 0.2 μl (0.06 U) of low-glucose Amyloglucosidase (Sigma, A7420-5MG) in 9.8 μl of PBST and incubated at 37 °C for 30 minutes; Glucose Oxidase (GO) reagent (25 μl) from the Glucose (GO) Assay Kit (Sigma, GAGO20) was added to determine total glucose + glycogen. Another 2 μl homogenate was mixed with 25 μl GO reagent alone (without the Amyloglucosidase) to determine free glucose. To determine trehalose contents, 1 μl supernatant was incubated with 0.5 μl Trehalase from porcine kidney (Sigma, T8778) with 9.5 μl PBST and incubated at 37°C overnight. Glucose Oxidase (GO) reagent (25 μl) was then added and incubated at 37 °C to determine trehalose + free glucose. Free glucose readings were subtracted from measurements of glycogen + glucose and trehalose + glucose to obtain readings for glycogen and trehalose, respectively. All absorbances were measured using an EnSight multimode plate reader (PerkinElmer), and values were normalized to protein content.

To measure the lipid and glucose contents of excreta, 3 days after temperature shift, flies were incubated with 0.5% BPB-supplemented fly medium for an additional day for prefeeding. Then, groups of five flies were transferred to 2-ml tubes containing BPB food cast into the lid of the tube for an additional 24 hours at 30 °C. After carefully removing the animals and cutting of the lids, tubes were washed with 100 μl of PBST, gently agitated with fresh lids and read at 540 nm absorbance for normalization. To measure the lipid and glucose contents of excreta, 10 μl of extracts were used and quantified as described above.

### Locomotor assays

To analyse long-term locomotor activity of individual flies, we used the *Drosophila* activity monitor system (TriKinetics). Individual 4-day old flies of the appropriate genotype were housed in 65-mm-long glass tubes containing food (5% sucrose and 2% agar in water) in one end and a cotton plug in the other, which were assembled into the monitoring system. The experiments were run in a behavioral incubator at 25 °C with 50-60% humidity and a 12-hour:12-hour light-dark circadian rythm. The flies were allowed to acclimate for 14 hours prior to data acquisition. The activity of individual flies was then measured as the number of beam crossings per 10 minutes, which was calculated using pySolo software in combination with an in-house MATLAB script. The stereotypic early activity period (EAP) and late activity period (LAP) were used to assess significant changes in locomotor activity.

We also analyzed short-term parameters of locomotor activity using a custom-built activity monitor system. Flies (n=10-16) were isolated individually in Plexiglas chambers, and allowed to acclimate for 2 hours. Flies were then recorded at the same time for 5 minutes using a high-resolution digital camera. Following an experimental cycle, parameters such as activity traces, total distance traveled and the maximal response velocity for each individual fly was calculated in MATLAB.

### Histochemistry

Adult brains, muscles, and fat bodies were dissected in PBS and fixed in 4% PFA in PBS for 20 minutes. Next, tissues were washed twice in PBS, incubated with periodic acid solution (Merck #395B-1KT) for 5 min, and washed twice in PBS. Samples were then stained with Schiff’s reagent (Merck #395B-1KT) for 15 minutes, washed twice in PBS, and mounted in 50% glycerol in PBS. Images were acquired with an BX-51 microscope equipped with an digital camera (Olympus). Two independent experiments were conducted and gave the same results.

### Starvation resistance

Flies were raised at the permissive temperature (18 °C) until 24 hours after eclosion, after which they were transferred to the restrictive temperature (30 °C) to disinhibit GAL4. Three days after transgene activation at 30 °C, flies were transferred unto 1% non-nutritive agar and incubated at 30 °C. Each vial was then observed every 8 hours until all flies (n=213-288) were counted as dead.

### Ramsay fluid secretion assay

Secretion assays were performed as described previously^60^. Intact Malpighian tubules from 7-day-old female flies were dissected in AHL and isolated into a 10 μl drop of a 1:1 mixture of AHL and *Drosophila* saline. The tubules were left to secrete for approximately 30 minutes, with non-secreting tubules being replaced if necessary, to produce a set of 10-20 working tubules. A drop of secreted fluid was subsequently collected every 10 minutes, and the diameter was measured using an eyepiece graticule. The volume of each droplet was calculated as 4Πr^3^/3, where r is the radius of the droplet, and secretion rates were plotted against time. Secretion was measured under basal conditions in order to establish a steady rate of secretion, prior to stimulation with labeled or unlabeled Capa peptides, with an increase in fluid secretion rate being taken as an indication of a diuretic effect.

### Statistics

Numerical data are presented as means ± s.e.m. The normal (Gaussian) distribution of data were tested using D’Agostino-Pearsen omnibus normality test. A one-tailed Student’s *t*-test was used to evaluate the significance of the results between two samples. For multiple comparisons tests, a one-way ANOVA followed by Tukey’s multiple comparisons of means was applied. Survival curves were assessed by the log-rank (Mantel–Cox) test and conducted for each pairwise comparison. Significance levels of P<0.05 (*), P<0.01 (**), P<0.001 (***) and P<0.0001 (****) were used in all tests. The statistical tests were performed using the data analysis program GraphPad Prism 8.0 (GraphPad Software Inc., CA, USA).

## AUTHOR CONTRIBUTIONS

T.K., S.T. and K.A.H. designed and conceptualized the study. T.K., S.T., M.T.N, S.N. and K.A.H. performed the experiments and analyzed the data. K.A.H. wrote the manuscript with input from S.T. All authors reviewed and edited the manuscript. K.A.H. produced the figures, directed the project and provided the funding.

## ACKNOWLEDGEMENTS

We are grateful to Manfred Frasch, Achim Paululat, Diego Manuel and Dr. Jae Park, for the generous sharing of resources, as well as to Michael Texada for giving critical comments to the manuscript. Camilla Trang Vo and Christina Papamichail are thanked for helping with pupal weight and developmental timing quantifications. This work was supported by funding from the Villum Foundation (15365) and Danish Council for Independent research Natural Sciences (9064-00009B) to KAH. Additional funding was given by UKRI BBSRC (BB/P008097/1) to SD, JATD and ST as well as by Novo Nordisk Foundation (16OC0021270) to K.R.

## SUPPLEMENTAL INFORMATION

**Figure S1.**
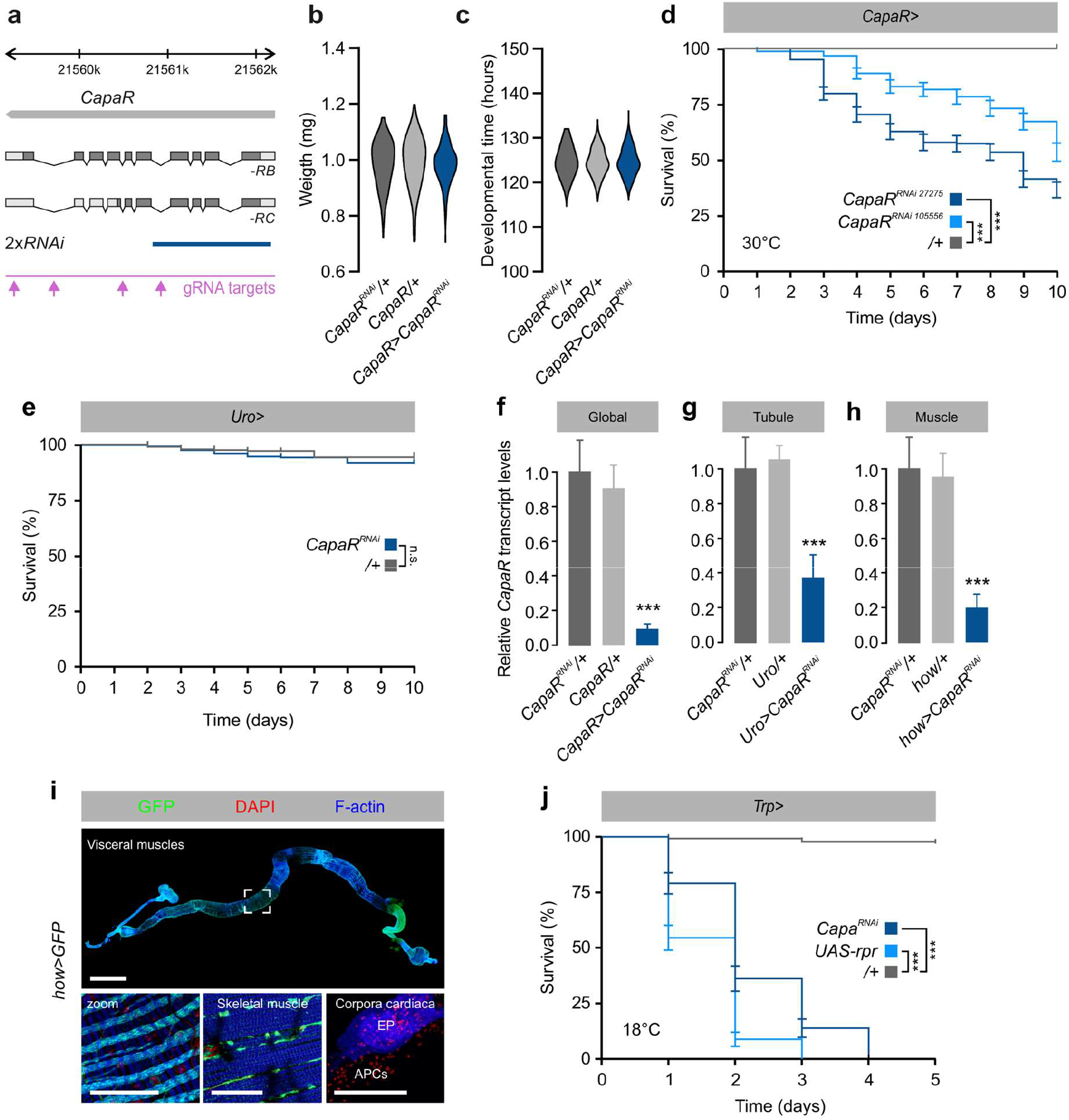
Extra-renal Capa/CapaR signaling is essential for adult fly viability. **a.** Exon map of the *CapaR* gene indicating the regions targeted by RNAi (blue; in-house generated RNAi line) and CRISPR/Cas9-mediated (magenta) genome editing techniques. **b.** Pupal weight of global *CapaR* knockdown flies (*CapaR>CapaR^RNAi^;* n=134) is not significantly different compared to *CapaR>* (n=111) and *CapaR^RNAi^/+* (n=140) parental controls (one-way ANOVA). **c.** Developmental time of global *CapaR* knockdown flies (*CapaR>CapaR^RNAi^;* n=129) is not significantly different compared to *CapaR>* (n=107) and *CapaR^RNAi^/+* (n=123) parental controls (one-way ANOVA). **d.** Two independent *CapaR^RNAi^* lines, *CapaR^RNAi 27276^* (n=183) and *CapaR^RNAi 105556^* (n=151), show the same mortality phenotype relative to control (n=151) confirming the specificity of the RNAi-effect (log-rank test). **e.** Knockdown of *CapaR* in tubule principal cells (n= 287) did not induce significant changes in fly survival compared to control (log-rank test; n=231). **f.-h.** *CapaR* transcript abundance normalized to *RpL32* following RNAi-mediated *CapaR* knockdown **d.** globally, **e.** in Malpighian tubules, and **f.** in muscles (one-way ANOVA; n=5). **i.** Analyzing *how>mCD8::GFP* (green) confirm GAL4-driven expression in visceral and skeletal muscles, but not the CC, of adult *Drosophila.* Scale bar 75 μm. **j.** Knocking down *Capa* expression (n=72) or completely ablating (n=70) the Capa-producing neurons by ectopic expression of *rpr*, phenocopies the mortality of *CapaR* silencing in adult *Drosophila* relative to control (log-rank test; n=133).

**Figure S2.**
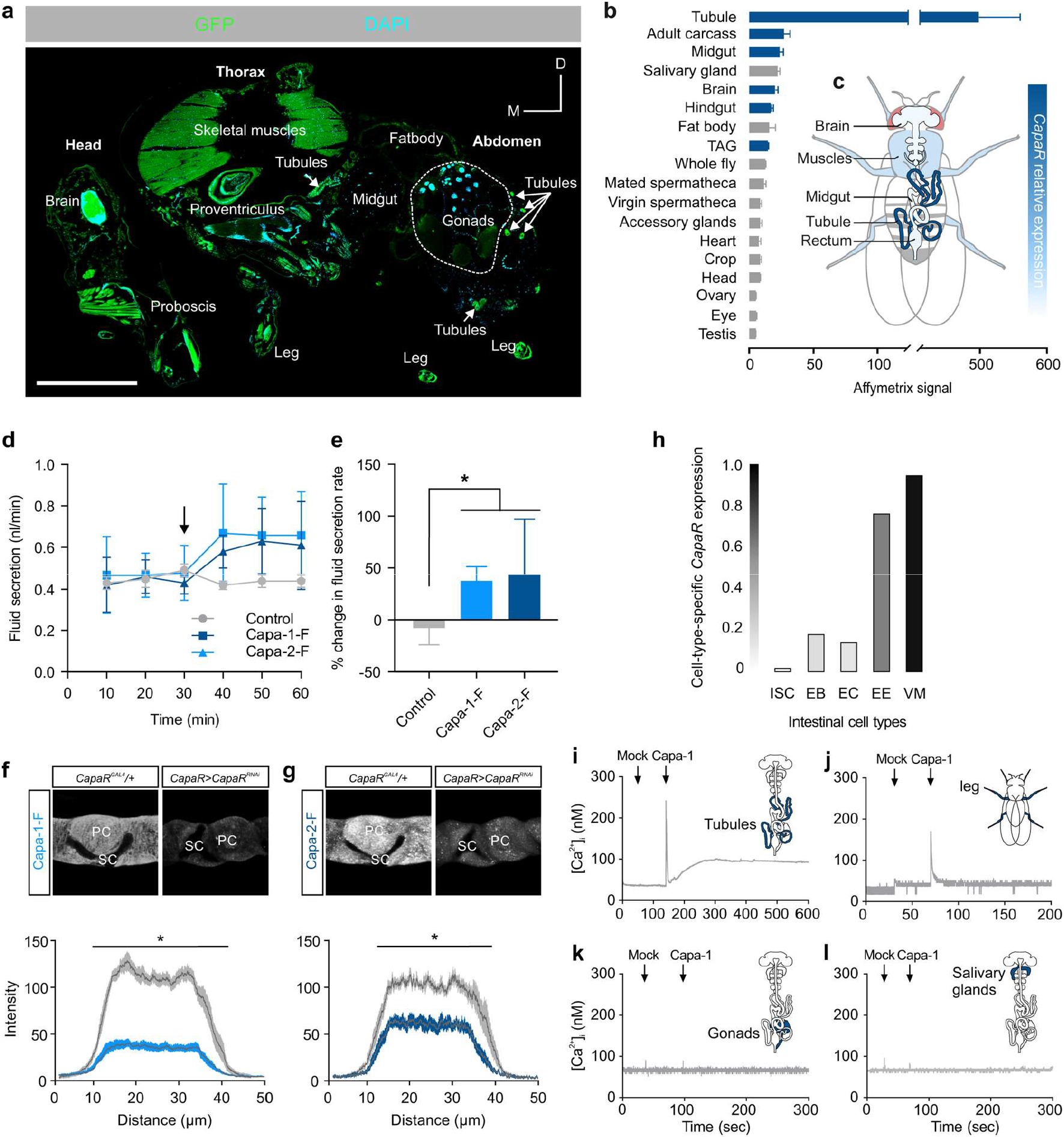
Spatial expression analysis and functional validation of CapaR. **a.** Medial paraffin section of adult fly carrying *CapaR>* and *UAS-mCD8::GFP* showing GFP expression in different tissues, including the tubules, brain, somatic musculature, and gut. **b.** Mean normalized Affymetrix signal ± s.e.m showing *CapaR* spatial expression across major adult tissues. **c.** Relative *CapaR* expression superimposed on an anatomical map of adult *Drosophila.* **d.-e.** Validation of fluorophore-coupled Capa-1 (Capa-1-F) and Capa-2 (Capa-2-F) peptides. **d.** Fluid secretion assays show significant functional stimulation of renal tissues by Capa-1-F and Capa-2-F peptides demonstrating biological activity. **e.** Percent change in fluid secretion rate following addition of Capa-1-F or Capa-2-F ligands (one-way ANOVA; n= 7). **f.** Knockdown of *CapaR* expression significantly reduces fluorescent intensity in principal cells compared to parental controls following Capa-1-F and Capa-2-F application (one-way ANOVA; n=5). **h.** Relative expression of *CapaR* in the different cell types of the intestine as realized by FACS sorted cell populations. Intestinal stem cells, ISC; Enteroblasts, EB; Enterocytes, EC; Enteroendocrine cells, EEs; Visceral muscles, VM. Data from: flygutseq.buchonlab.com^62^. **i.-l.** Functional validation of tissue-specific Capa signaling using *in vivo* calcium-reporter technology. Stereotypic biphasic increases in cytosolic [Ca^2+^]_i_ was detected upon Capa-1 application (10^−7^ M) in **i.** tubules, and **j.** legs, but not in **k.** testes or **l.** salivary glands.

**Figure S3.**
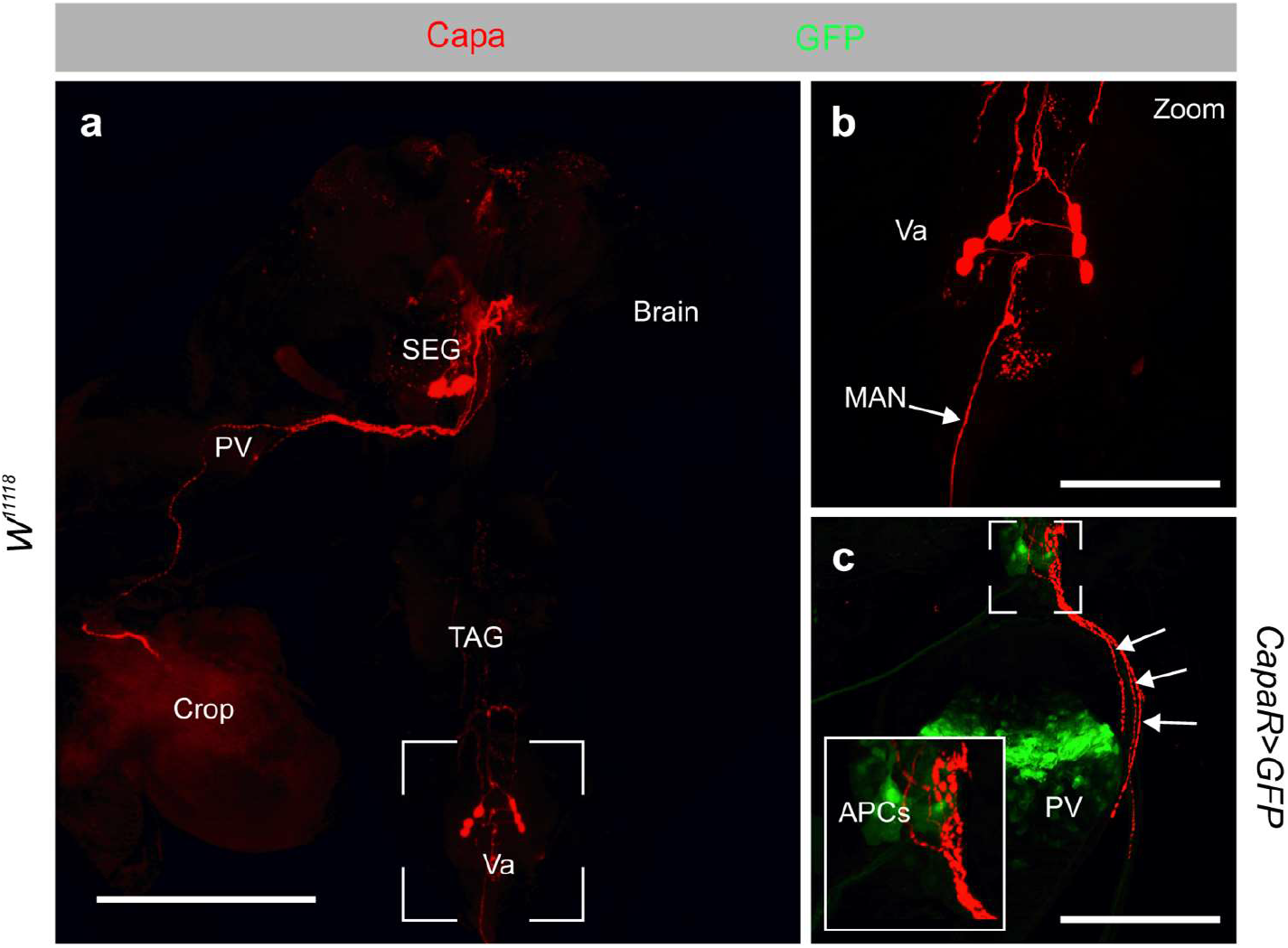
Neuroanatomy of Capa^+^ neurons. **a.** Immunoreactivity of Capa precursor antibody (red) identifies one pair of SEG neurons and three pairs of Va Capa^+^ neurons. Scale bar = 200 μm. **b.** The Va neurons release Capa peptides into circulation via the median abdominal neuron (MAN). Scale bar = 100 μm. **c.** The axons from the SEG neurons innervate the AKH-producing neuroendocrine cells (APCs) and proventriculus (PV), which also shows *CapaR>mCD8::GFP* expression (green) in muscle cells. Scale bar = 100 μm.

**Figure S4.**
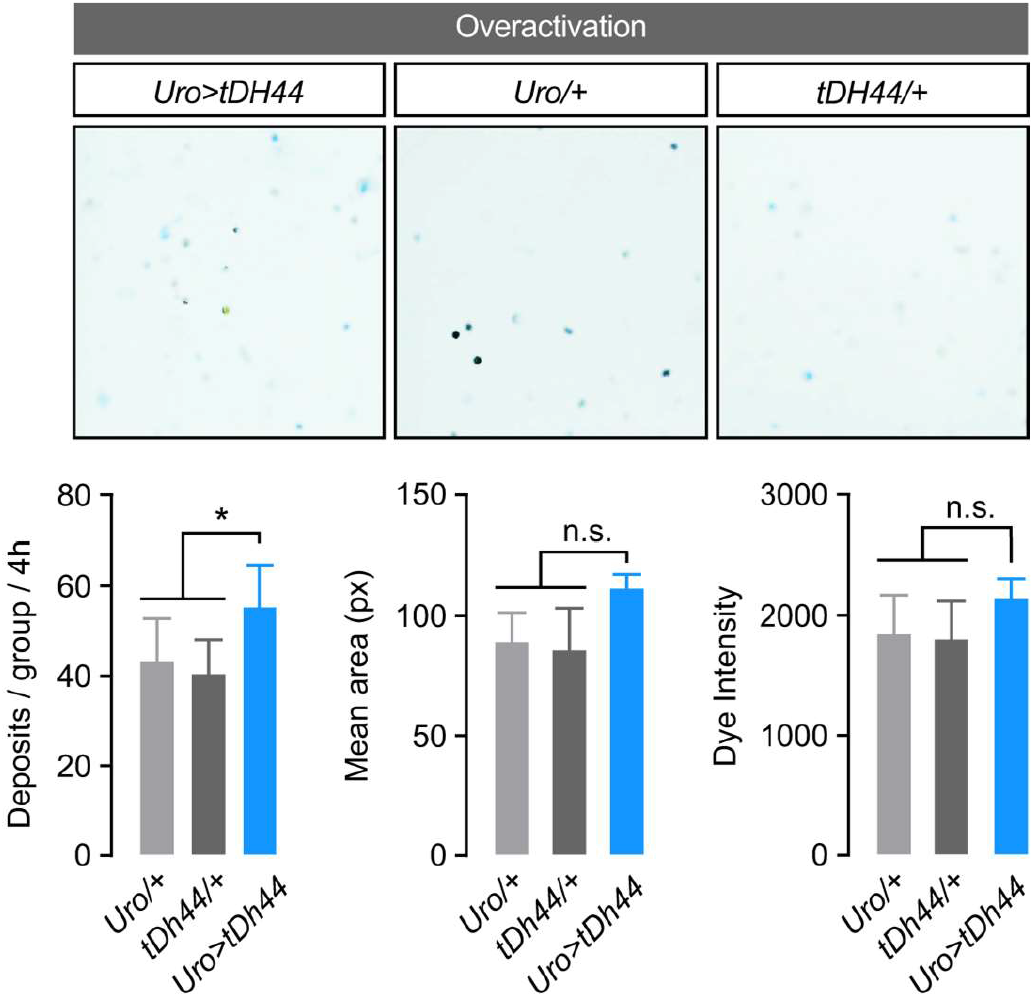
Functional stimulation of renal tubules increase waste excretion. Fecal output profiles following artificial activation of renal secretion by ectopic expression of a membrane-tethered version of diuretic hormone 44 (tDH44) show that these flies only produce more, but not larger of more dilute, deposits compared to control (one-way ANOVA; n=5-7).

**Figure S5.**
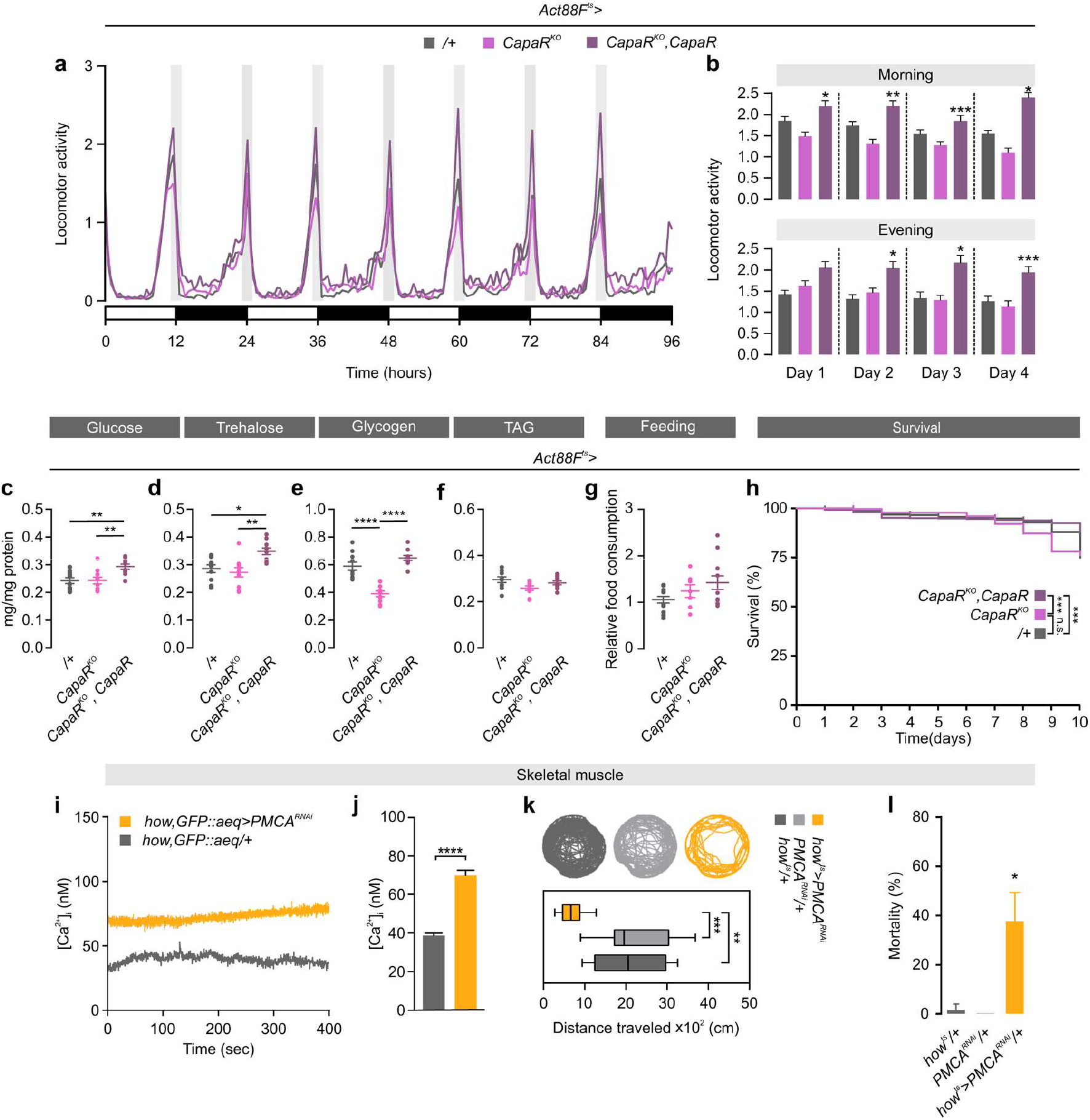
Physiological effects of eliminating Capa signaling in skeletal muscles. **a.** Locomotor activity of individual control (+/; n=32), skeletal muscle-specific *CapaR* knockout (*Act88F^ts^>CapaR^KO^;* n=30) and rescue (*Act88F^ts^>CapaR^KO^,CapaR;* n=31) flies exposed to 12-hour:12-hour, light-dark (LD) cycles over 96 hours. **b.** Morning and evening activity peaks measured during the experimental period (one-way ANOVA). **c.-f.** Quantification of c. glucose, d. trehalose, e. glycogen and f. TAG levels in flies with the different genotypes (one-way ANOVA; n=10). **g.** Relative food intake (one-way ANOVA n=10). **h.** Kaplan-Meier survival curves of control (+/; n=508), knockout (*Act88F^ts^>CapaR^KO^*; n=594) and rescue (*Act88F^ts^>CapaR^KO^,CapaR;* n=677) flies (log-rank test). **i.-j.** Quantification of [Ca^2+^]_i_ following *PMCA* knockdown in skeletal muscles (*how, GFP::aeq>PMCA^RNAi^*) relative to control (*how,GFP::aeq/+;* Student’s *t*-test; n=3). **k.** Representative activity traces of video-tracked individual flies with targeted *PMCA* knockdown in muscles (*how^ts^>PMCA^RNAi^;* n=10-16) relative to control flies, including quantification of distance travelled (one-way ANOVA). **l.** Knockdown of *PMCA* in muscles (*how^ts^>PMCA^RNAi^*) causes significant fly mortality (one-way-ANOVA; n=60).

**Figure S6.**
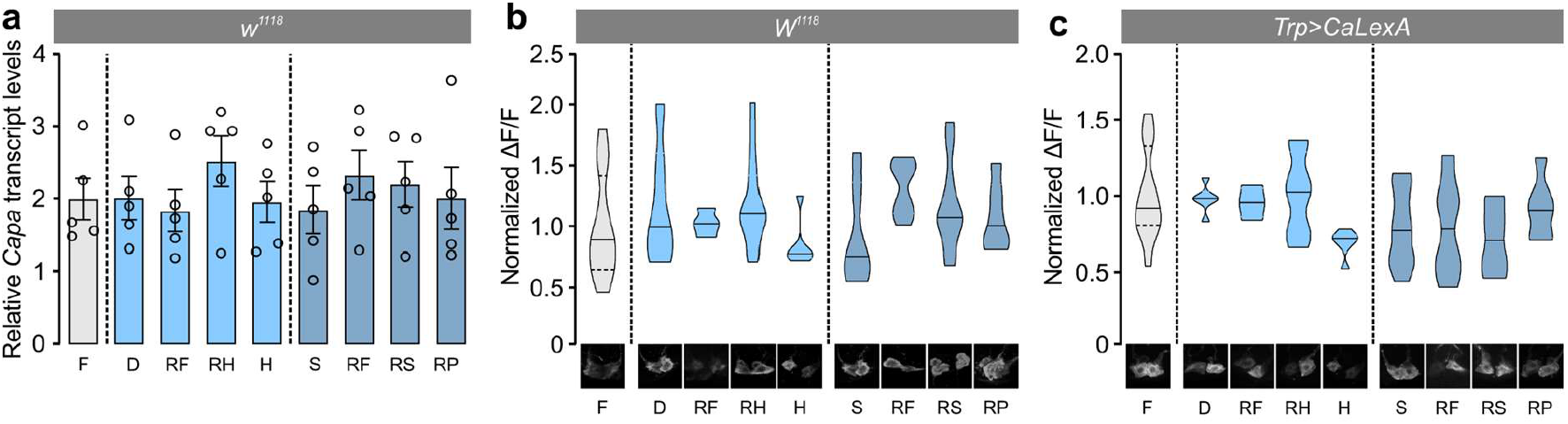
Capa^+^ SEG neuron activity is unaffected by water or nutrient stress. **a.** Transcript levels of *Capa* relative to *RpL3* in the SEG neurons (head samples) from flies exposed to different environmental conditions (one-way ANOVA; n=5). **b.** Violin plots of immunofluorescence quantifications of intracellular Capa precursor levels (n=6-16) and **c.** CaLexA induced GFP expression in Capa-producing SEG neurons (n=4-12) from flies exposed to different environmental conditions (one-way ANOVA). F, fed; D, desiccated; RF, refed; RH, rehydrated; H, hydrated; S, starved; RS, refed sugar; RP, refed protein.

**Table S1.**
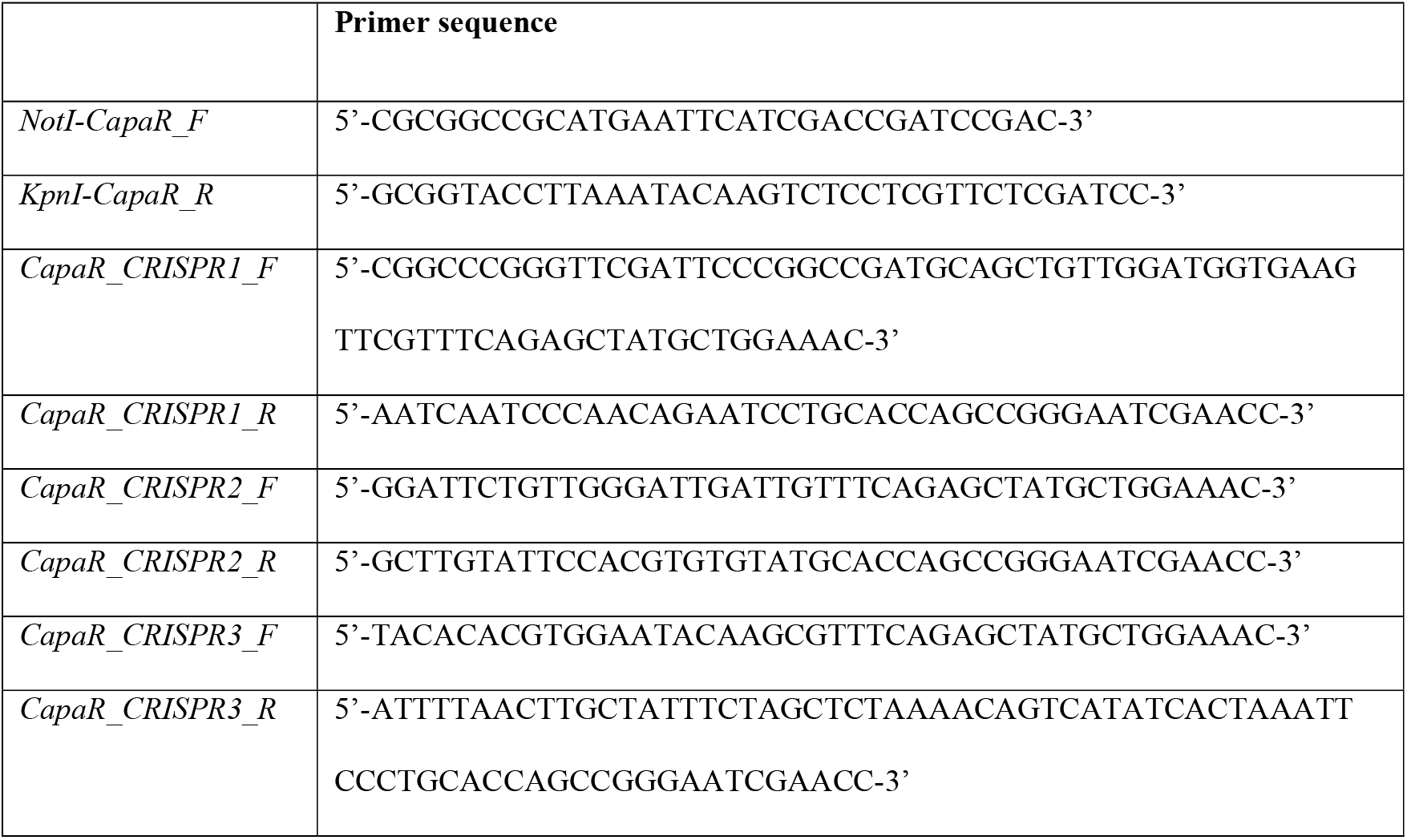
Primer sequences used in *Drosophila* transgenesis.

**Table S2.**
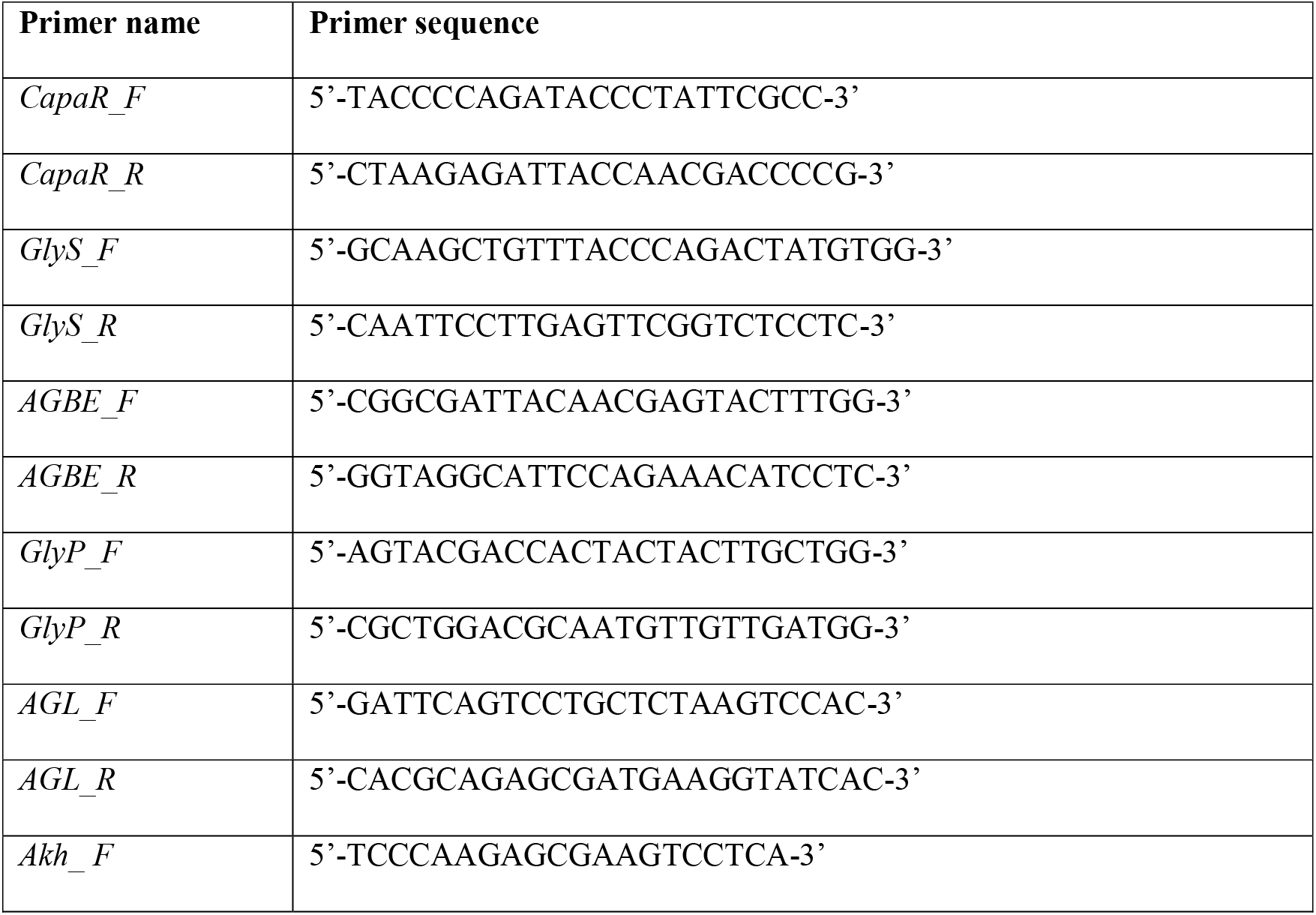
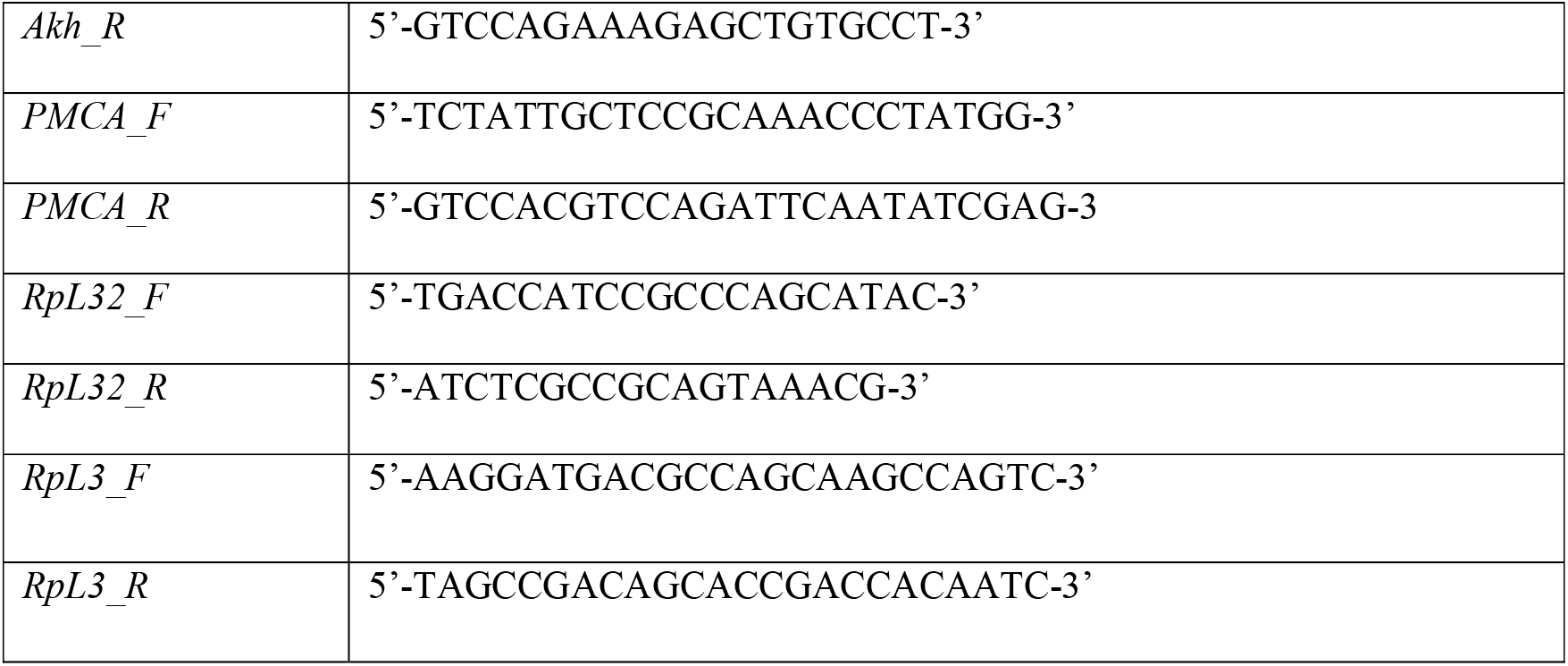
Quantitative RT-PCR primers used.

## Notes

### Competing Interest Statement

The authors have declared no competing interest.

